# TCR Signal Strength and Eomes Coordinate CD8^+^ T Cell Fate and Antitumor Immunity in Hepatocellular Carcinoma

**DOI:** 10.64898/2026.02.10.705135

**Authors:** Hacene Dreidi, Abdulkader Azouz, Sébastien Denanglaire, Solange Dejolier, Faustine Bouffioux, Marie Le Moine, Soren Temara, Lune Cassart, Muriel Nguyen, Lionel Larbanoix, Stanislas Goriely, Fabienne Andris

## Abstract

Hepatocellular carcinoma (HCC) poses a persistent barrier to effective immunotherapy, in part due to the emergence of exhausted CD8^+^ T cells. Using a hydrodynamic, autochthonous HCC model based on Sleeping Beauty-mediated delivery of oncogenic drivers and defined OVA-derived antigens, we could manipulate antigen affinity under physiological priming conditions. With this system, we examined how TCR signal strength and the transcription factor Eomesodermin (Eomes) shape CD8^+^ T cell fate in vivo. High-affinity stimulation supported robust effector differentiation and the formation of tissue-resident memory T cells (Trm), resulting in early tumor control. In contrast, low-affinity stimulation induced elevated Eomes expression and favored rapid transition toward an exhausted (Tex) phenotype. Genetic repression of Eomes enhanced Trm differentiation in high-affinity CD8^+^ T cells and delayed tumor relapse, whereas ectopic Eomes expression accelerated exhaustion and led to earlier tumor recurrence. Under low-affinity conditions, Eomes deficiency increased the Trm-to-Tex ratio but remained insufficient to restore effective antitumor immunity, indicating that weak TCR engagement imposes constraints that cannot be overcome by Eomes loss alone. These findings reveal that the strength of TCR engagement shapes CD8^+^ T cell fate in HCC, while Eomes selectively biases this process toward exhaustion at the expense of effector and resident memory programs, ultimately influencing the durability of tumor control.

**Synopsis:** This study reveals that the strength of T-cell receptor signaling fundamentally steers whether CD8^+^ T cells in liver cancer become durable tumor-fighting cells or slip into exhaustion, pinpointing Eomes as a key factor that pushes cells toward dysfunction. By uncovering how antigen affinity and Eomes jointly shape antitumor immunity, the work offers new conceptual and therapeutic avenues for improving the persistence and effectiveness of immunotherapies in hepatocellular carcinoma.

## Introduction

Hepatocellular carcinoma (HCC) is the most common primary liver malignancy and a leading cause of cancer-related mortality worldwide. Despite advances in systemic therapies, including immune checkpoint inhibitors, prognosis remains poor, with only approximately 20% of patients showing clinical benefit, largely due to tumor immune evasion and the complexity of the tumor microenvironment ^1–4^. As an inflammation-driven cancer, HCC is shaped by both the quality and quantity of immune cell infiltration, as well as their dynamic interactions within the tumor microenvironment, all of which critically influence the immunotherapy outcomes ^4,5^. Recent studies have underscored the pivotal role of tumor-infiltrating lymphocytes, particularly CD8^+^ T cells, in orchestrating the immune response against HCC. These cells develop through various functional states, including effector T cells (Teff), exhausted T cells (Tex), and tissue-resident memory T cells (Trm). The balance between these states strongly correlates with prognosis and response to immunotherapy ^6–8^.

T cell exhaustion arises during chronic antigen stimulation, such as persistent infections and cancer and is characterized by sustained expression of inhibitory receptors and progressive loss of effector functions ^9–12^. In HCC, exhausted T cells contribute to immune dysfunction and are associated with poor prognosis ^13^. Although transcriptomic signatures of exhaustion have been developed to predict survival and guide immunotherapy ^6,8,14,15^, the heterogeneity and functional plasticity of Tex cells remain incompletely understood.

Trm cells differ from circulating memory cells by their expression of retention markers (CD69, CD49a) and local effector functions such as secretion of IFN-γ and granzyme B ^16–18^. Liver Trm support early disease resolution in non-alcoholic steatohepatitis (NASH) and fibrosis ^19^ and a higher Trm-to-Tex ratio predicts better outcomes under PD-1 blockade in HCC ^13^. However, Trm can acquire exhaustion-like features, including PD-1 expression and functional impairment, likely driven by chronic antigen exposure and a suppressive tissue milieu ^20–22^. In line with this, we recently showed that PD-1 signaling curtails the oligoclonal expansion of liver Trm through Eomes-dependent regulation ^23^.

Eomes, a T-box transcription factor, plays a pivotal role in CD8^+^ T cells differentiation. Eomes supports effector and memory formation ^24–26^, promotes survival through IL-2/IL-15 receptor signaling ^25,27,28^, and participates in early lineage bifurcation by favoring the development of circulating memory T cells while restraining residency programs ^29^. Conversely, elevated expression of Eomes is a hallmark of exhausted CD8^+^ T cells in chronic viral infections and in cancer, where it is associated with increased inhibitory receptor expression (TIM-3, PD-1) and reduced cytokine production ^30–33^. Thus, Eomes exerts context-dependent, and sometimes opposing effects on CD8^+^ T cell differentiation.

The strength and duration of T cell receptor (TCR) signaling adds another layer of regulation, profoundly influencing T cell fate decisions. Persistent, strong TCR stimulation drives differentiation toward an exhausted phenotype, whereas transient stimulation or intermediate antigen levels favor the development of effector and memory T cells ^34^. In acute infections, Eomes is selectively induced by weak activating signals and promotes the survival of low-affinity clones, thereby broadening the memory pool ^25^. How TCR affinity and Eomes interact to shape T cell fate in solid tumors and particularly in the antigen-rich, tolerogenic liver microenvironment remains poorly understood.

In this study, we investigated how Eomes and antigen affinity jointly regulate CD8^+^ T cell differentiation in an autochthonous, antigen-defined HCC model. We show that strong TCR signaling is required for the generation of effector and tissue-resident T cells, and for early tumor control. Eomes repression limits T cell exhaustion and promotes Trm differentiation, thereby prolonging tumor control in high-affinity CD8^+^ T cells. Under low-affinity stimulation, characterized by higher Eomes expression, its repression increases the Trm-to-Tex ratio, but fails to restore effective antitumor function. Together, these findings reveal how TCR strength and Eomes activity converge to shape CD8^+^ T cell fate in HCC.

## Results

### High-affinity tumor-specific T cells control early hepatocarcinoma progression

To investigate CD8^+^ T cell response to HCC, we implemented a versatile model of autochthonous hepatocarcinoma based on hydrodynamic delivery of oncogene-encoding plasmids and defined OVA-derived antigens via the Sleeping Beauty (SB) transposon system for genomic integration ^35^. We selected a combination of myristoylated Akt1 (myrAkt1), mutated RAS (NrasG12V) and a dominant-negative form of TP53 (R246S), together with shRNA targeting endogenous *Trp53* transcripts (plasmids NrasG12V + Akt + Tp53+ sh3’Tp53), which elicited a strong OVA-specific immune response, detected by in vivo pentamer staining ^35^ (Fig S1A, B).

The SB plasmid also encoded Luciferase, enabling non-invasive monitoring of tumor burden. Tumors were induced in CD3ε-deficient mice, allowing unrestricted tumor growth in the absence of host T cells. Three weeks later, transfer of naive RAG1^−/−^ OT-I cells in OVA-HCC bearing mice triggered a rapid decrease in bioluminescence (days 6-7 post OT-I cell transfer), followed by tumor re-growth 3-4 weeks later, demonstrating transient antitumoral response but limited long-term tumor control (Figure S2A, B).

Control mice injected with OVA transposon alone (OVA-Liver) showed no evidence of tumor development as confirmed by macroscopic inspection and liver weight (Fig S2C, D). By day 7, similar numbers of OT-I cells were recovered from the liver of OVA-Liver and OVA-HCC mice, and both groups upregulated exhaustion markers (PD1, TOX) compared to untreated mice receiving naive OT-I cells. Although HCC-bearing mice showed a lower percentage of PD1^+^TOX^+^, compared to OVA-liver counterparts, they expressed higher proportion of terminally exhausted T cells (TCF7^low^ TIM3^high^), and fewer polyfunctional cytokine-producing cells, indicating more advanced exhaustion driven by antigenic stimulation in a tumoral context (Fig S2 E-M).

To assess how antigen affinity shapes T cells differentiation in HCC, we compared OT1 responses to transposons encoding high-affinity SIINFEKL (N4) peptide or altered peptide ligands of reduced affinities: Q4 (SIIQFEKL) and T4 (SIITFEKL), following the hierarchy N4 > Q4 > T4 (Fig 1A). Despite comparable baseline tumor burden across groups, only high-affinity stimulation (N4) induced robust tumor regression by day 7 (Fig 1B, C and Fig S3A). Very few T cells were recovered from the T4-HCC group, precluding further analysis. In contrast, N4-HCC and Q4-HCC groups yielded similar numbers of infiltrating OT-I cells (Fig 1D). Both groups contained similar numbers of PD1^+^ TOX^+^ (Tex-like) T cells, but their relative abundance was higher in Q4-HCC mice (Fig 1E-F). However, consistent with previous studies ^36^, t-SNE analysis revealed clear phenotypic divergence between N4- and Q4-primed OT1 cells, with high affinity priming preferentially driving effector-like cells (CX3CR1, KLRG1, TIM3) and acquisition of tissue-residency markers (CXCR6, CD49a) (Fig S3B). We further confirmed increased frequencies and numbers of TCF1^low^ CX3CR1^+^ (Effector-like) and CXCR6^+^ CD49a^+^ (Trm-like) OT-I cells in N4-HCC mice (Fig 1G-J). Similar observations were made in the spleen, although the recovery of OT-I cells was lower in this organ, with very few cells exhibiting Trm features (Fig S4). Given the prognostic value of Trm/Tex ratio in HCC ^13^, we compared this index across groups. N4-HCC mice consistently displayed higher Trm/Tex ratios (Fig 1K). Functionally, N4-primed OT-I cells produced increased amounts of granzyme B, IFNγ, and TNFα, whereas Q4-primed produced more IL-2 (Fig 1L-O, Fig S3C-F), further reflecting divergent functional states despite comparable expansion.

**Figure 1.**
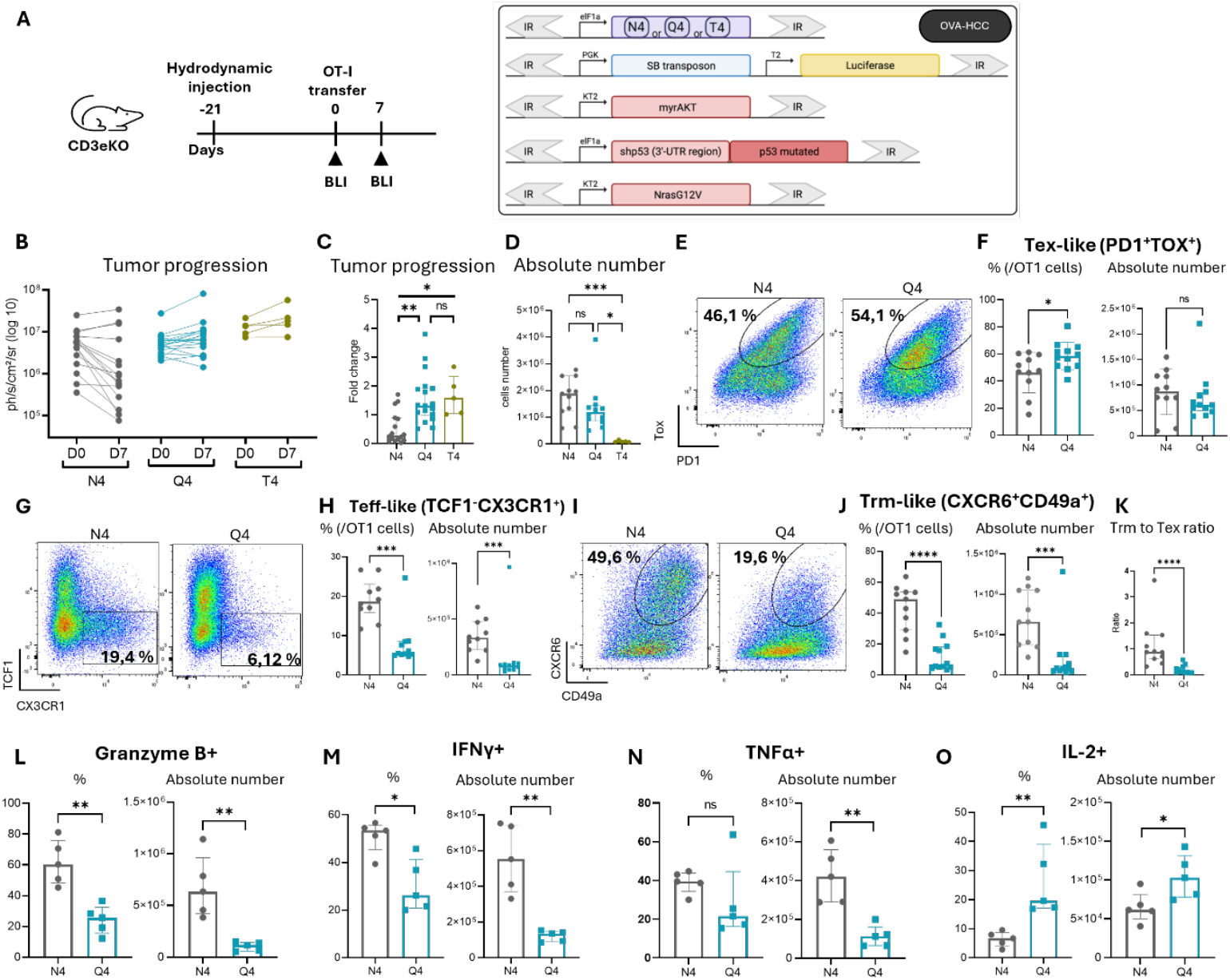
Antigen affinity governs the balance between effector, tissue-resident, and exhausted OT-I CD8^+^ T cells in HCC. CD3ε^−^/^−^ mice were hydrodynamically injected with HCC-inducing plasmids combined with altered OVA peptide transposons, and OT-I cells were transferred 3 weeks later. **(A)** Experimental design to investigate differences in TCR affinity in the HCC model (left panel) and list of plasmids co-injected hydrodynamically (right panel). N4, Q4, T4 refer to the high-affinity SIINFEKL (N4), intermediate-affinity SIIQFEKL (Q4), and low-affinity SIITFEKL (T4) Ova peptide encoding plasmids, respectively. **(B)** Bioluminescence imaging (BLI) quantification of tumor burden (photon flux, ph/s/cm^2^/sr) before (day 0) and one week after OT-I cell transfer (day 7). Individual mouse values are shown for each HCC groups, as indicated. **(C)** Fold-change in BLI signal (day 7/day 0) for mice from panel A. **(D)** Absolute numbers of OT-I CD8^+^ T cells recovered from tumoral livers at day 7 in the N4, Q4 and T4 groups. **(E-J)** Representative flow cytometry plots (E, G, I) and frequencies and absolute numbers (F, H, J; left and right panels) of PD-1/TOX (E, F), CX3CR1/TCF1 (G, H), and CD49a/CXCR6 (I, J) on gated OT-I CD8^+^ T cells from N4 and Q4 tumoral livers at day 7. **(K)** Ratio of tissue-resident memory (Trm) to exhausted (Tex) OT-I CD8^+^ T cells in tumoral livers at day 7 for N4 and Q4 mice. **(L-0)** Frequencies and absolute numbers of Granzyme B^+^ (L), IFN-γ^+^ (M), TNF-α^+^ (N), and IL-2^+^ **(O)** OT-I CD8^+^ T cells at day 7. Individual mice are shown for each experimental group; n= 5 to 12 (pool of 2 independent experiments (B-K) or representative experiment out of 2 (L-O). Statistical significance was determined using the Kruskal-Wallis test (B-C) or the Mann-Whitney test (E-O), ns = non-significant, *p<0.05, **p<0.01, ***p<0.001, ***p<0.0001. *See also Figures S3-S5*

Together, these findings indicate that continuous but subthreshold TCR signaling favors the acquisinon of an exhausted phenotype over Teff and Trm features. In contrast, high-affinity TCR engagement enables both effective antitumor response and the generation of effector and tissue-resident memory CD8+ T cells.

### Eomes promotes transition to Tex and inhibits Trm differentiation in HCC

We next examined the expression of transcription factors associated with T cell exhaustion in response to low-affinity antigen stimulation. In Q4-HCC mice, a large fraction of liver-infiltrating OT-1 cells strongly express Eomes, with a similar but weaker trend in the spleen (Fig 2A, B). In contrast, only a small fraction of OT-I cells expressed Eomes in N4-HCC mice. Notably, these Eomes^+^ cells displayed reduced CD49a and CX3CR1 expression, compared with their Eomes^-^ counterparts (Fig S5A, B), consistent with the established role Eomes as a negative regulator of Trm and Teff cell development ^28,29^. Moreover, Eomes^+^ cells also exhibited increased TCF1 expression in both N4-HCC and Q4-HCC mice (Fig S5C). To directly assess the functional impact of Eomes on antitumor activity, we first compared the response of WT and *Eomes*^*Flox/Flox*^. CD4-Cre (Eomes-KO) OT-I cells in the full-length OVA-HCC model (Fig 2C). Although both WT and Eomes-KO OT-I cells effectively suppressed tumor growth by day 7, Eomes-KO recipients showed markedly reduced tumor rebound at days 14 and 21 (Fig 2D and S6). At day 7, the total number of liver-infiltrating T cells was higher in the Eomes-KO group, but this difference diminished at later time points (Fig 2E).

**Figure 2.**
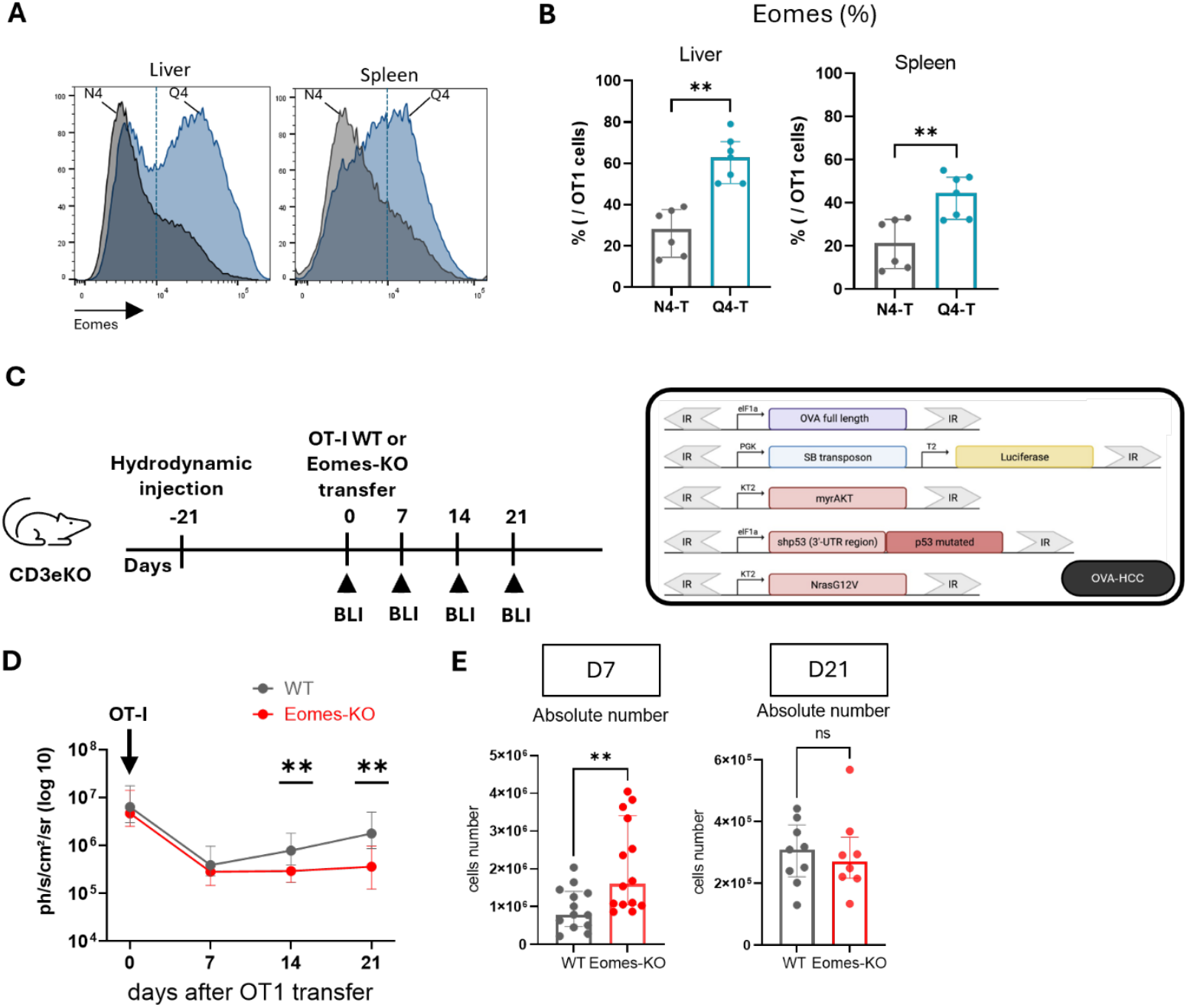
Eomes depletion improves tumor control. **(A-B)** Representative histograms (A) and percentages of Eomes expressing cells (B) in OT-I CD8^+^ T cells from liver and spleen of N4 and Q4 mice at day 7. **(C)** Experimental design to study the depletion of Eomes in the HCC model. **(D)** Longitudinal bioluminescence imaging (BLI) quantifying tumor burden (photon flux, ph/s/cm^2^/sr) before (day 0) and following OT-I cell transfer for 3 weeks. **(E)** Absolute numbers of OT-I cells recovered from the liver 7- and 21-days post-transfer. Data are shown as median with interquartile range and are representative of 2 experiments with n= 6-7 mice per group (B) or are pooled from 2 to 3 experiments with n= 8-13 (D, E). Statistical significance was determined using the Mann-Whitney test, ns = non-significant *p<0.05, **p<0.01, ***p<0.001, ****p<0.0001. Mann-Whitney test was corrected for multiple comparisons using Holm-Šídák method.

To explore how Eomes shapes T cell differentiation during tumor relapse, we performed scRNAseq analysis on OT-I cells recovered from HCC-bearing livers at day 21. Of a total of 11025 OT-I cells, unsupervised clustering identified 12 clusters grouped into 8 transcriptionally defined subsets (Fig 3A). Based on gene expression and defined signatures, we identified central memory, effector memory, Trm, cycling, ISAG^hi^, and various exhausted populations. Four stem-like/Tcm clusters (Tcm1–4), characterized by hierarchical expression of *Tcf7, Klf3*, and *Lef1*, were collectively defined as Tcm subsets. Three exhaustion-associated clusters were identified: Tex (*Tox, Pdcd1, Eomes, Bhlhe40*), precursor cells (Tpex) (*Tox, Pdcd1, Tcf7, Myb, Slamf6*), and Tex-eff (*Tox, Pdcd1, Tnfrsf9, Xcl1, Crtam, Nr4a2/3, Cd160*) (Fig. 3A, D and Fig. S7). WT OT-I cells were predominantly enriched for Tex and Tpex subsets, whereas Eomes-KO OT-I cells displayed increased proportions of Trm, Tem and Tcm subsets (Fig 3B). Differential gene expression within Tcm, Tex and cycling subsets revealed that *Eomes* deficiency enhanced expression of genes and signature related to residency (*P2rx7, Cxcr6, Itga1, Itga4, Itgb1, Itgb7, Itgae*) across multiple lineages (Fig 3C, D and Fig S8A-C, F), indicating that the absence of Eomes not only expands “bona-fide” Trm cell clusters but also induces residency-associated programs in additional subsets. In contrast, consistently with low expression of Eomes in this subset, Tem subsets displayed few transcriptional changes between WT and Eomes-KO cells, indicating that in the absence of Eomes, their increased frequency was not accompanied by substantial reprograming (Fig S8D, E). Slingshot analysis using Tcm1 cells as the root population revealed 5 distinct developmental trajectories, 4 of which initially progress through the Tcm1-4 clusters. Lineages 1 and 3 are predominantly observed in the WT group. In these trajectories, Tcm4 cells either directly progress to Tex (lineage 3) or first acquire Trm1 features, then transition through Tpex state, ultimately exhibiting a Tex-eff phenotype (lineage 1). Lineages 4 and 5 are primarily found in the Eomes-KO group. Lineage 4 cells directly progress from Tcm4 to Tem, while lineage 5 diverges earlier (from Tcm2 state) and transition to ISAG^hi^, culminating in Trm2. Lineage 2 is shared between both groups and goes directly from Tcm4 to the cycling subset. These findings indicate that Eomes constrains developmental pathways that would otherwise support effector or resident memory differentiation (Fig 3E).

**Figure 3.**
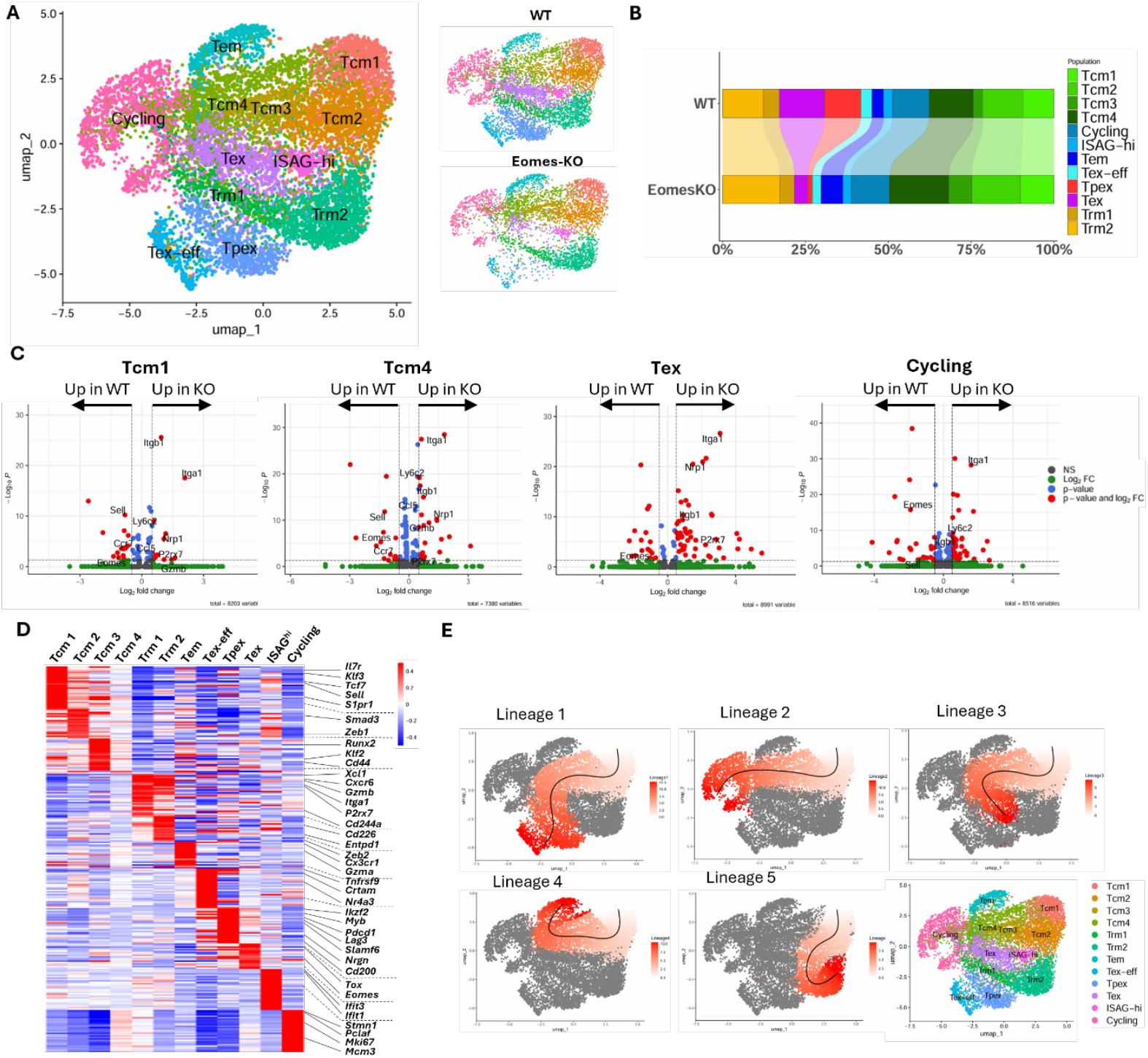
Eomes-KO T cells that develop in HCC-bearing mice acquire a transcriptional program associated with residency. **(A)** Uniform manifold approximation and projection (UMAP) plots of OT-I CD8^+^ cells from the livers of WT and Eomes-KO mice and annotated in 8 distinct populations, according to their patterns of gene expression, represented as global UMAP (left) and split in WT and Eomes-KO conditions (right). **(B)** Frequencies of each annotated cluster (WT and Eomes-KO groups). **(C)** Volcano plot comparisons of selected clusters from WT and Eomes-KO OT-I cells recovered from HCC livers. Each dot represents one gene. Red dots correspond to p value < 0,01 and fold change > 0,5. **(D)** Heat map showing relative expression of selected genes in each cluster with representative genes indicated on the right. **(E)** Trajectory analysis using Slingshot. The five lineages are shown separately on the UMAP. *See also Figures S7 and S8*

FACS analysis performed on day 7 (a timing when both groups exhibit the same tumor burden) indicated early divergence in cell fate. Eomes deficiency markedly increased CD49a expressing T cell subsets, including cells co-expressing PD-1, TOX, CXCR6, and CD69, suggestive of an intermediate transitory state between precursors of Trm and Tex cells (Fig 4A).

**Figure 4.**
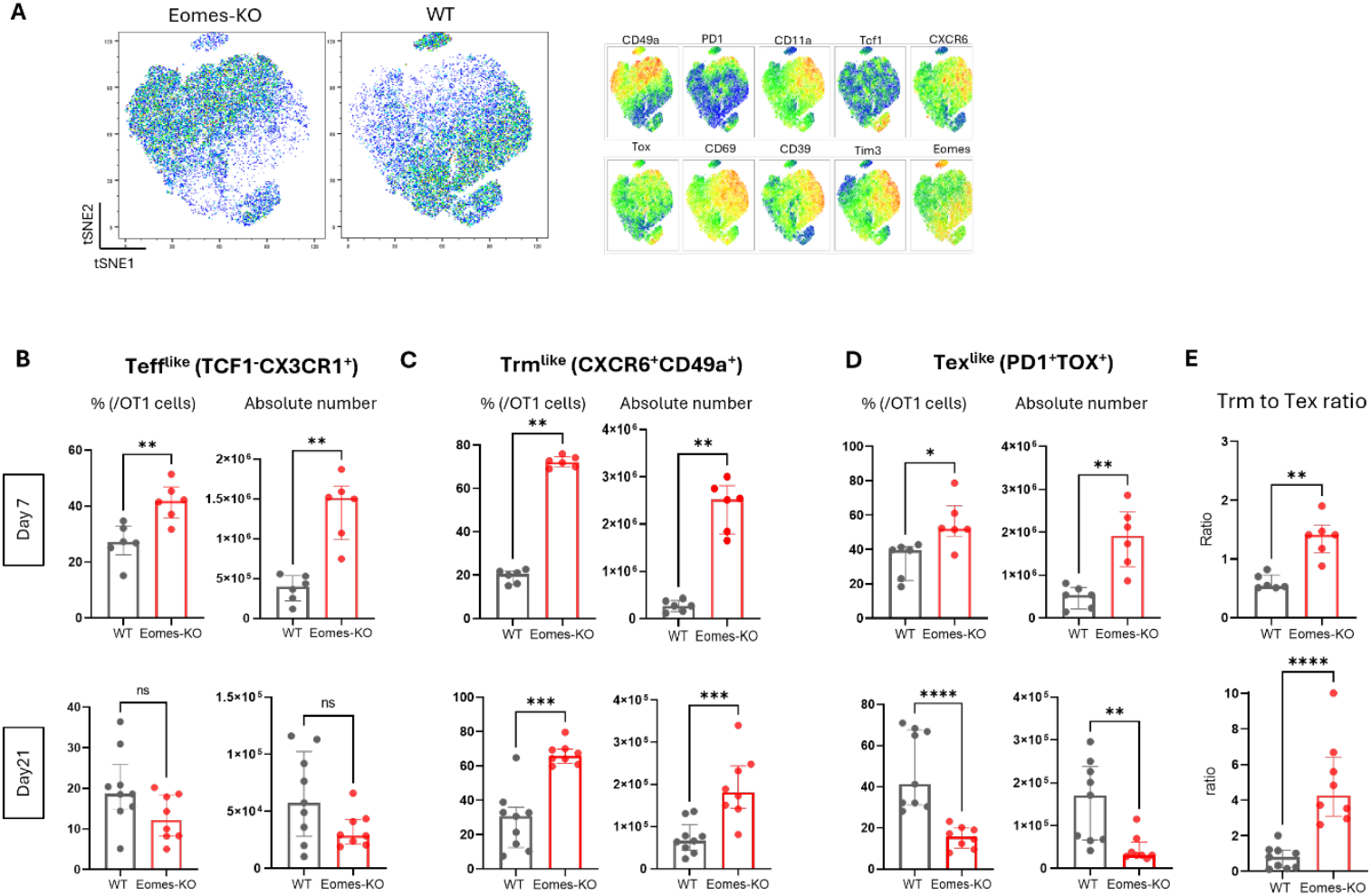
Eomes promotes transition towards Tex subset and inhibits Trm differentiation in a HCC. CD3ε^−^/^−^ mice were hydrodynamically injected with OVA-HCC-inducing plasmids, and WT or Eomes-KO OT-I cells were transferred 3 weeks later. **(A)** t-SNE analysis of WT and Eomes-KO OT-I cells, showing expression of selected markers. **(B-D)** Proportion (right panels) and absolute numbers (left panels) of effector-like cells (B), tissue-resident memory-like cells (C) and exhausted-like cells (D) subpopulations at day 7 (upper row) and day 21 (lower row). Markers used for population identification are indicated on top of histograms. **(E)** Ratio of Trm-like to Tex-like cells in the liver at day 7 (upper panel) and 21 (lower panel). Data are shown as median with interquartile range and are representative of 2-3 experiments (n= 5 to 9 mice per group), each dot represents a single mouse. Statistical significance was determined using the Mann-Whitney test, ns = non-significant, *p<0.05, **p<0.01, ***p<0.001, ****p<0.0001. (B) Mann-Whitney test was corrected for multiple comparisons using Holm-Šídák method.

At both day 7 and day 21, Eomes-KO groups exhibited expansion of Trm-like cells (CD49a^+^ CXCR6^+^) (Fig 4C and Fig S9A), whereas Tex-like (PD1^+^TOX^+^) cells initially increased at day 7 but later contracted (Fig 4D and Fig S9B). Accordingly, the Trm/Tex ratio progressively increased in the absence of Eomes (Fig 4E, compare upper and lower panels). Eomes-KO group also displayed increased proportions and absolute numbers of effector-like cells (TCF1^-^ CX3CR1^+^) on day 7 (Fig 4B, upper panels). By day 21, comparable total numbers of this cell subset were recovered in both groups of mice, but a substantial fraction of WT late Teff-like expressed TOX, while Eomes-KO cells did not (Fig S9C-E). This observation suggests that part of this subset represents Tex-eff cells.

Given the link between cytotoxic potential and CX3CR1 expression ^30,37^, as well as the association of KLRG1 with short-lived effector cells, we evaluated the expression of these markers. Eomes deficiency increased the frequency of CXC3CR1^+^ cells while reducing KLRG1^+^ and CXCR3^+^KLRG1^+^ dual positive cells - recently associated with tissue circulating long-lived effector cells ^38^ (Fig S10, lower panels). Functionally, Eomes KO OT-1 cells produced more Granzyme B, IFNγ and, to a lesser extent, TNFα and IL-2, further supporting enhanced T cell functionality in this group (Fig S11).

In summary, Eomes promotes commitment toward exhausted states and restrains both effector and tissue-resident memory differentiation. Loss of Eomes enhances CD8^+^ T cell functionality, skews differentiation toward Trm and effector lineages, and improves the durability of antitumor control in HCC.

### Eomes deficiency enhances Trm development in low-affinity HCC transposon but does not restore effective tumor control

To evaluate whether Eomes depletion could rescue Trm differentiation and improve tumor control under low affinity antigen stimulation, we repeated the experiments using the Q4-HCC transposon model (Fig 5A). On day 7, a time point when only high-affinity (N4-primed) OT-I cells control tumor growth (Fig 1B, C), the absence of Eomes did not restore tumor regression in Q4-HCC mice (Fig 5B). Absence of early tumor regression in this context precluded further longitudinal studies for ethical reasons. We did not observe significant differences in the total number of OT-I cells recovered from the livers at day 7 (Fig 5C). Nevertheless, Eomes-KO OT-I cells displayed clear phenotypic shifts in response to low-affinity stimulation. Specifically, Eomes KO cells were enriched in TCF1^lo^ subset expressing CD49a, CXCR6, CD39, CD69, TIM3 and, to a lower extent, CX3CR1 (Fig S12). Proportion and absolute numbers of Tex-like cells decreased in the absence of Eomes (Fig 5F), whereas frequency (but not total numbers) of both Trm-like (CD49a^+^ CXCR6^+^) and Teff-like (TCF1^-^ CX3CR1^+^) cells increased in the Eomes-KO group (Fig 5D, E), leading to higher Trm/Tex ratio in KO mice (Fig 5G).

**Figure 5.**
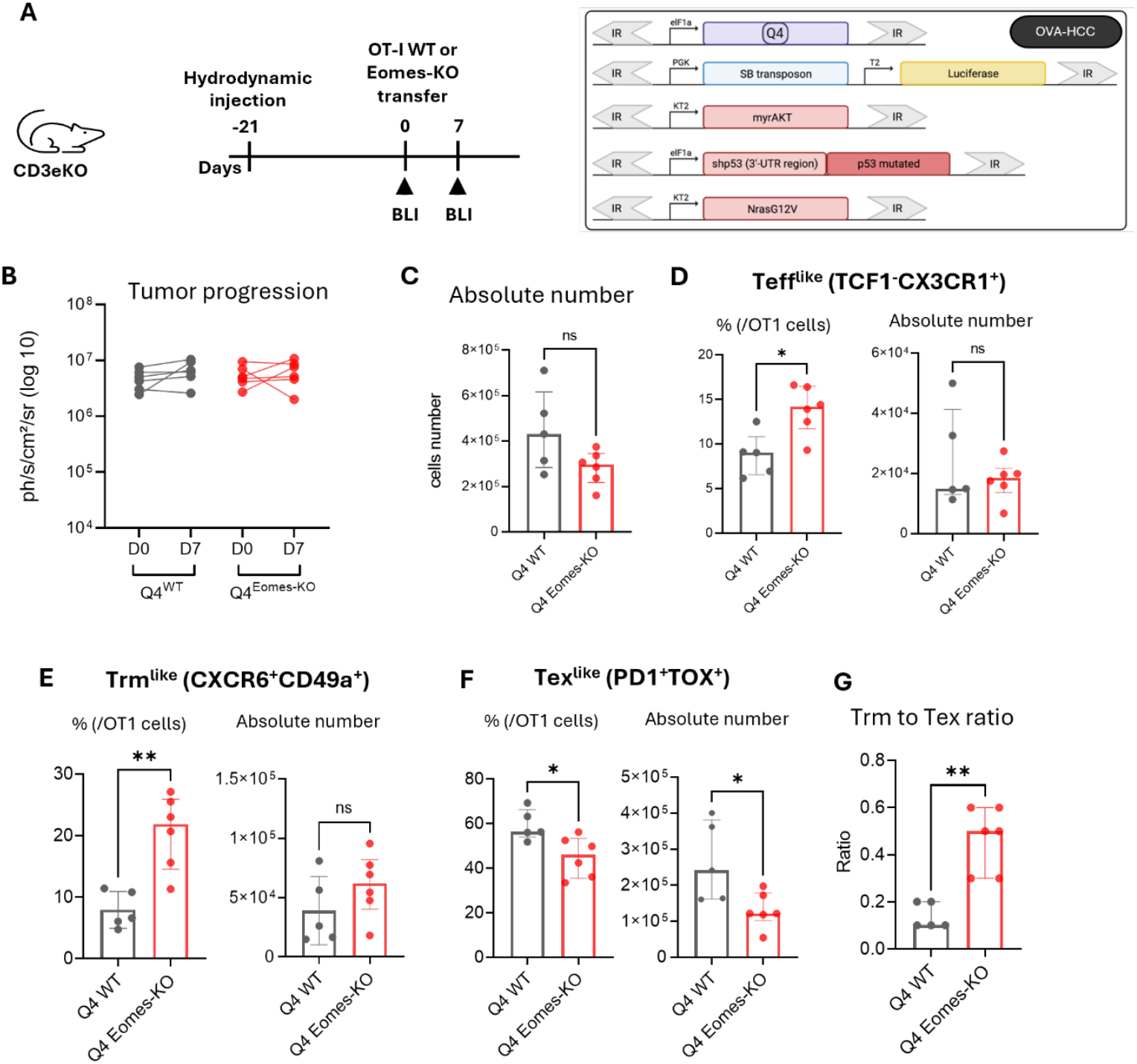
Eomes deficiency enhances Trm development but does not restore tumor control in low-affinity HCC model. (A)Experimental design to study the depletion of Eomes in a low-affinity HCC model. **(B)** BLI quantification of tumor burden before (day 0) and 7 days after transfer of WT and Eomes-KO OT-I cells in Q4 peptide-expressing HCC recipient mice. **(C)** Absolute number of OT-I cells recovered after 7 days in each group. **(D-F)** Frequencies (left panels) and absolute numbers (right panels) of Teff-like (D), Trm-like (E) and Tex-like (F) cells in WT and Eomes-KO OT-I recipient mice. **(G)** Ratio of Trm-like to Tex-like cells in each group of mice. Data are shown as median with interquartile range (n= 5-6 mice per group), each dot represents a mouse. Statistical significance was determined using the Mann-Whitney test, ns= non-significant, *p<0.05, **p<0.01.

Thus, even under low-affinity antigen stimulation, the absence of Eomes promotes Trm and Teff development at the expense of Tex, mirroring observation in the high-affinity model. However, these shifts fail to restore effective antitumor immunity, indicating that the intrinsic limitations imposed by weak TCR engagement cannot be overcome by Eomes’ repression alone.

### Transgenic Eomes expression restrains T cell expansion and enforces exhaustion in HCC

To further define the role of Eomes in T cell differentiation in the context of HCC, we took advantage of a transgenic mouse strain expressing Eomes under the control of hCD2 regulatory elements (named *Eomes*^*Tg*/Tg 23,24^), crossed with *Rag1*^*−/−*^ OT-I mice (Fig 6A). In these mice, OT-I cells express Eomes under steady-state conditions but at levels that remain in a physiological range and that does not result in spontaneous PD-1 nor TOX expression (Fig S13A, B). Ectopic Eomes expression does not preclude initial (day 1 and 2) OT-I T cell proliferation after OVA stimulation in vitro but leads to overall decreased cell survival and/or proliferation at later time points (Fig S13C).

**Figure 6.**
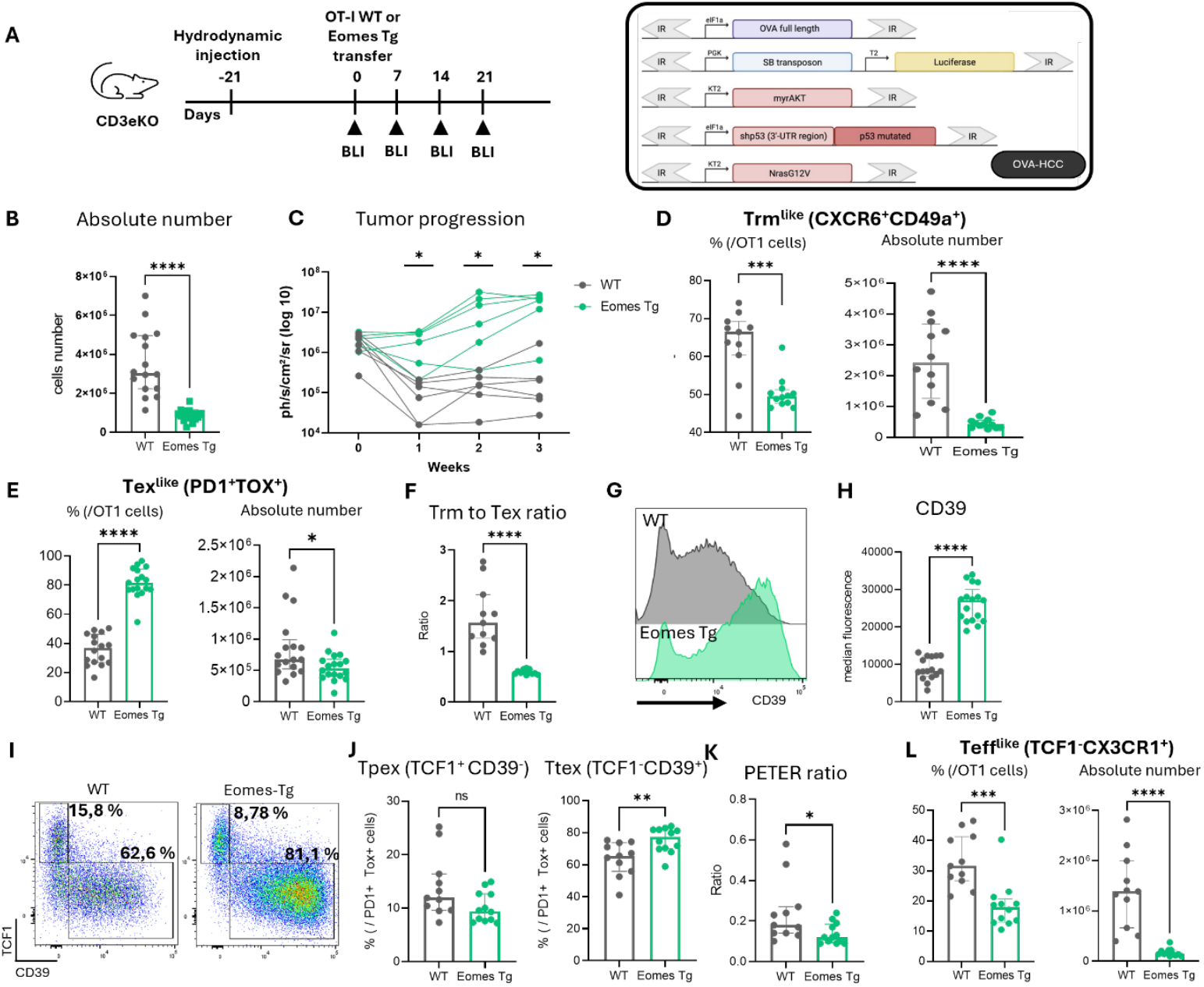
Transgenic expression of Eomes in OT-I cells restrains their immune control of OVA-HCC. (A)Experimental design to study transgenic Eomes expression in HCC model. **(B)** Absolute numbers of WT and Eomes-Tg OT-I cells recovered on day 7 after transfer in OVA-HCC recipient mice. **(C)** BLI quantification of tumor burden before (day 0) and following OT-I cell transfer for 3 weeks. **(D-E)** Frequencies (left panels) and absolute numbers (right panels) of Trm-like (D) and Tex-like (E) in cells from (B). **(F)** Ratio of Trm-like to Tex-like cells in each group of mice. **(G, H)** Representative histograms (G) and median fluorescence intensity (MFI) (H) of CD39 expression in WT and Eomes-Tg OT-I cells. **(I, J)** FACS profiles (I) and frequencies (J) of TCF1^+^CD39^-^ precursor exhausted cells (Tpex) and TCF1^-^TOX^+^ terminally exhausted cells (Ttex) among PD1^+^TOX^+^ OT-I CD8^+^. **(K)** Peter ratio calculated from (J). **(L)** Frequencies (left panels) and absolute numbers (right panels) of Teff-like cells. Data are shown as median with interquartile range and are pooled from 2-3 experiments (with n= 11 to 17), each dot represents a single mouse. Statistical significance was determined using the Mann-Whitney test, ns= non-significant, *p<0.05, **p<0.01, ***p<0.001, ****p<0.0001. (B) Mann-Whitney test was corrected for multiple comparisons using Holm-Šídák method.

Lower numbers of OT-I.*Eomes*^Tg/Tg^ were recovered from HCC livers after adoptive transfer compared to their WT counterparts (Fig 6B). Despite this reduced expansion, a large fraction of HCC-bearing mice exhibited early tumor regression (day 7) following transfer of OT-I.*Eomes*^Tg/Tg^ (Fig S14A). However, these mice developed accelerated tumor relapse, compared to WT OT-I recipients, demonstrating defective long-term control of the tumor when Eomes expression is enforced (Fig 6C).

In agreement with the inhibitory role of Eomes on residency program (Fig 3B, Fig 4C), enforced Eomes expression led to decreased proportion and total numbers of Trm-like cells (CD49a^+^ CXCR6^+^) (Fig 6D, see also Fig S14B, C for representative 2-D dot plots and unsupervised t-SNE analysis). Conversely, Increased proportion of Tex-like PD-1^+^TOX^+^ cells developed among *Eomes*^Tg/Tg^ OT-I cells (Fig 6E and S14E). Yet, the total number of Tex-like cells was slightly reduced in these mice, reflecting the poor overall T cell recovery (Fig 6B). Moreover, the Trm/Tex ratio was markedly decreased in *Eomes*^Tg/Tg^ OT-I cells, confirming global bias toward exhaustion (Fig 6F).

We next examined markers associated with advanced exhaustion. The ectonucleotidase CD39, a hallmark of terminal CD8^+^ T cell exhaustion in both mice and humans ^39–41^, was upregulated in *Eomes*^Tg/Tg^ OT-I cells (Fig 6G, H). Analysis of TCF1 and CD39 expression revealed a higher proportion of Ttex-like TCF1^low^CD39^high^ cells in PD1^+^TOX^+^ cells from *Eomes*^Tg/Tg^ recipient mice (Fig 6I, J). Calculation of the PETER index - the ratio of Tpex-to-Ttex ^42^ - confirmed a shift toward advanced exhaustion upon ectopic Eomes expression (Fig 6K). Moreover, effector-like (TCF1^-^ CX3CR1^+^) T cells were significantly decreased in *Eomes*^Tg/Tg^ HCC mice (Fig 6L and S14D).

Collectively, these experiments strongly support the notion that Eomes promotes exhaustion upon both high- and low-affinity antigenic stimulation, at the expense of residency and effector profiles and restrains durable antitumoral response even when early control is achieved.

## Discussion

In the present study, we demonstrate that both TCR affinity and Eomes expression critically shape the fate and antitumor activity of CD8 T cells in HCC. We show that submaximal TCR stimulation induces high Eomes expression in OVA-specific tumor-infiltrating T cells, consistent with prior reports in acute CMV and *Listeria monocytogenes* infections ^25^. In our autochthonous HCC model, low-affinity stimulation drives differentiation toward exhausted (Tex-like) phenotypes, whereas high-affinity stimulation promotes robust effector (Teff) and tissue-resident memory (Trm) differentiation, improved cytokine production, and superior tumor control. These findings establish a direct link between TCR signal strength and Eomes-regulated lineage bifurcation in tumor-infiltrating CD8^+^ T cells.

Although strong TCR stimulation is often associated with terminal CD8^+^ T cell exhaustion in chronic infections ^43^, the relationship between TCR affinity and dysfunctional CD8^+^ T cell states in cancer is more complex. Recent data from a Lewis lung carcinoma model indicate that optimal TCR engagement promotes the development of Tpex cells in tumor-draining lymph nodes and supports their intra-tumoral persistence, whereas reduced TCR stimulation accelerates the terminal differentiation of Tex cells ^44^. Conversely, in the MCA205 fibrosarcoma model, high TCR signal strength was required to establish a dysfunction-associated molecular program in tumor-specific T cells ^34^. The reasons for these discrepancies remain unclear but may relate to differences in tumor models, tissue context, or antigen expression levels across studies. Importantly, these studies did not evaluate the development of effector or resident memory T cell subsets, limiting direct comparison. Nevertheless, our findings support the hypothesis that low affinity signaling in the HCC environment may be sufficient to drive exhaustion, whereas strong TCR signals preferentially support protective Teff and Trm programs.

Modulation of Eomes profoundly impacted CD8^+^ T cell fate and tumor progression under high-affinity stimulation. Eomes deficiency enhanced effector and Trm differentiation and conferred prolonged control of HCC, whereas Eomes constitutive expression promoted exhaustion and accelerated tumor relapse.

Across multiple studies, Eomes emerges as a critical transcription factor shaping CD8^+^ T cell fate and antitumor function, with both beneficial and detrimental effects reported. In chronic lymphocytic leukemia or under OX-40/CTLA-4 costimulatory therapy, Eomes is required for the generation of potent effector CD8^+^ T cells and for sustaining antitumor activity ^45,46^. In contrast, other studies report that elevated Eomes expression is linked to T cell exhaustion, loss of co-activating receptors, and impaired tumor control in solid cancers. In these contexts, cells harboring high Eomes expression exhibit a dysfunctional phenotype, limiting immunotherapy efficacy ^31,47^. Moreover, although Eomes promotes the generation of central memory T cells in peripheral lymphoid organs, it restricts long-term stemness in intra-tumoral CD8^+^ T cells ^48^. Our results strongly support the negative role for Eomes in the immune control of HCC. Our scRNAseq analysis revealed that Eomes deletion led to the expansion of Trm and Teff-like cells but also to an enrichment of stem-like subsets (e.g., Tcm4) that display a cell-cycling signature yet retain lower stemness compared with other Tcm states. This suggests a dual role for Eomes: maintaining stemness (beneficial for the renewal of tumor-specific T cells), while limiting their proliferation and transition toward effector and Trm cells, which mediate effective tumor clearance. Conversely, Eomes expression by effector cells may accelerate their progression to an exhausted state.

A dynamic crosstalk exists between the tumor and the immune system, and it is often challenging to disentangle the direct role of Eomes from the influence of tumor antigenic load, which may differ between experimental groups. By analyzing early time points, when tumor burden was comparable across groups, we demonstrate that Eomes-driven differentiation program precedes and likely contributes to subsequent tumor progression. Whether increased antigenic load in later stages further amplifies exhaustion in Eomes-deficient cells remains an open question.

Compared to genetic or subcutaneous tumor cell line models, hydrodynamic delivery of transposons encoding oncogenes and defined surrogate tumor antigens directly into hepatocytes enables tumor initiation in the native liver microenvironment, maintaining liver-specific immune architecture and allowing precise control over antigen affinity ^35^. Use of CD3ε-deficient hosts allowed uncontrolled tumor growth prior to adoptive transfer of OT-I cells, conceptually analogous to lymphodepletion in adoptive T cell therapies ^49^. In this experimental setting, the absence of CD4 T cells could accelerate HCC relapse, as recent data indicate that tumor-specific CD4 T cells can be recruited to overcome CD8 T cell dysfunction following adoptive cell transfer in a murine B16F10 melanoma model ^50^. Whether CD4 help modulates Eomes-dependent differentiation in HCC remains an important question for future work.

Finally, the absence of Eomes also promoted Trm and Teff development at the expense of Tex under low-affinity stimulation. However, it did not restore meaningful tumor control in Q4-HCC. This underscores the fact that weak TCR signaling imposes intrinsic constraints on antitumor CD8^+^ T cell responses that cannot be overcome by modulating Eomes alone. It would be important to evaluate the effect of anti-PD1 treatment in this setting. Collectively, our findings reveal that TCR signal strength and Eomes expression cooperatively dictate the balance between effector, resident memory, and exhausted CD8^+^ T cell fates in HCC. These results suggest that fine-tuning Eomes expression, together with adjustments in TCR signaling, may enhance the durability and the quality of CD8^+^ T cell responses in liver cancer.

## Material and methods

### Mice

CD3ε−/−mice under C57BL/6 background were purchased from Jackson Laboratory (Bar Harbor, USA). *Eomes*fl/fl mice under C57BL/6 background were purchased from the Jackson Lab (B6.129S1(Cg)-Eomestm1.1Bflu/J) and were crossed to CD4Cre+ mice obtained from the Jackson Lab (Tg(Cd4-cre)1Cwi/BfluJ) to generate CD4Cre^+^ *Eomes*fl/fl mice. Eomes transgenic mice (*Eomes*Tg/Tg) were generated as previously described ^24^. OT1αβ^+/+^Rag1^−/−^ *Eomes*Tg/Tg and OT1αβ^+/−^Rag1^−/−^ Eomes KO mice were generated by intercrossing OT1αβ^+/+^ (C57BL/6-Tg(TcraTcrb)1100Mjb/J) and Rag1^−/−^ (B6.Cg-Rag1tm1.1Cgn/J), both purchased from the Jackson Lab, with *Eomes*^Tg/Tg^ or CD4Cre+*Eomes*fl/fl mice. OT-I. Rag1^−^/^−^ littermates without Cre or transgene served as wild-type controls. Mice were bred and housed under specific pathogen-free conditions (SPF). Mice used for the experiments were males and females between 7 and 12 weeks of age. All animal work was conducted in compliance with and after approval by the institutional Animal Care and local committee for Animal Welfare from the Biopark ULB Charleroi.

### In vivo experiments

Hydrodynamic injections were performed according to standard procedures ^51^ with a mixture of plasmids encoding myroistylated Akt (pKT2/CLP-AKT), mutated Ras (pT3/EF1a-NRASG12V) and mutated tumor suppressor P53 gene (pT3/EF1a-TRP53-R246S-sh3’TRP53), combined with a plasmid encoding either the full-length Ovalbumin protein (pT3/EF1a–Ovalbumin-full-length) or one of three OVA epitope variants: N4 (pT3/EF1a–SIINFEKL), Q4 (pT3/EF1a– SIIQFEKL), or T4 (pT3/EF1a–SIITFEKL). Each injection also included a plasmid encoding the Sleeping Beauty transposase fused to luciferase (pT2/C-Luc//PGK-SB13) to facilitate genomic integration and enable bioluminescent tracking.

Tumor development was monitored noninvasively using bioluminescence imaging. This imaging was performed by means of a Photon Imager Optima (Biospace Lab) that dynamically counted the emitted photons for at least 25 min, under anesthesia (4% and 2% isoflurane for initiation and maintenance, respectively) and after subcutaneous administration of 150 mg/kg of D-luciferin in saline (Promega). Image analysis was performed with M3Vision software (Biospace Lab). Regions of interest (ROIs) were drawn on the mice abdomen, and signal intensities were quantified individually for a time lapse of 5 min corresponding to the maximum signal intensity plateau. Analysis of light emissions from luciferase-expressing cells enabled real-time tracking of tumor growth and localization over time.

All plasmids were administered at a dose of 5000 ng per mouse, with the total injected DNA not exceeding 25,000 ng. Three weeks after hydrodynamic injection, mice received an adoptive transfer of 1 to 2 million OT-I cells intravenously.

### Cell preparation

For all experiments, organs were processed under sterile conditions. The tumor-bearing liver is flushed by injecting 5–10 mL of PBS into the portal vein to remove blood, which causes the liver to blanch. Liver is perfused with a solution containing DNase I (Roche, 10104159001) and Liberase (Roche, 05401020001) to enzymatically digest liver tissue prior to mechanical dissociation. The digestion mix is injected throughout the hepatic lobes, and the liver is incubated at 37°C for 30 minutes.

After incubation, the reaction is quenched by adding approximately 5 mL of RPMI 5% supplemented with 2 mM EDTA. The liver is then mechanically dissociated using a syringe plunger and filtered through a 70 μm cell strainer into a 50 mL Falcon tube. Additional RPMI 5% EDTA 2 mM may be added as needed to maximize tissue dissociation. The cell suspension is centrifuged at 1400 rpm for 10 minutes at 4°C. The supernatant is discarded, and the cell pellet is resuspended in 8 mL of red HBSS, then filtered into a new tube.

To isolate T lymphocytes, the cell pellet is directly resuspended in 10-20 mL of Percoll. Centrifugation is performed at 2000 rpm for 30 minutes at 20°C without brake. T cells are found in the pellet, while debris and tumor cells remaining in the supernatant are discarded. Red blood cells were lysed by ACK lysing buffer (Ammonium-Chloride-Potassium, home-made).

Spleens were mashed and red blood cells were lysed by ACK lysing buffer. The cell suspension was neutralized by adding RPMI, 5% FBS and filtered before further manipulation.

### Flow cytometry

Single-cell suspensions were stained with live/dead fixable near-IR dead cell stain kit (Thermo Fisher Scientific, L34976) or iFluor 860 maleimide (AAT Bioquest, 1408) to exclude dead cells, incubated with Fc receptor–blocking antibodies CD16/32 (clone 2.4G2, BioXcell, BE0307, RRID:AB_2736987) to block non-specific binding and stained with the following fluorochrome-conjugated antibodies for surface markers: CD45.1 (A20, 565212, RRID:AB_2722493) TCRβ (H57-597, 560657, RRID:AB_1727575), CD8α (53-6.7, 563068, RRID:AB_2687548), CD69 (H1.2F3, 612793, RRID:AB_2870120), CD38 (90/CD38, 562768, RRID:AB_2737781), CD39 (Duha59, 143805, RRID:AB_2563393), KLRG1 (2F1, 565393, RRID:AB_2739216) Tigit (1G9, 744213, RRID:AB_2742062), TIM-3 (5D12/TIM-3, 747621, RRID:AB_2744187), PD-1 (J43, 562523, RRID:AB_2737634), CD3 (17A2, 565642, RRID:AB_2739318), CX3CR1 (Z8-50, 567820, RRID:AB_2916747), CXCR6 (SA051D1, 151109, RRID: AB_2616760), CD49a (Ha31/8, 740519, RRID:AB_2740235), H2-K^b^_SIINFEKL_-PE pentamer (F093-2B, PRO-Immune). For detecting intranuclear proteins, cells were fixed and permeabilized for 30 min with eBioscience Foxp3/Transcription Factor Staining Buffer Set (Thermo Fisher Scientific, 00-5523-00) before staining with the following antibodies: Eomes (Dan11mag, 53-4875-82, RRID:AB_10854265), TCF1 (S33-966, 564217, RRID:AB_2687845) and TOX (TXRX10, 50-6502-82, RRID:AB_2574265) from Thermo Fisher Scientific.

To evaluate the functional status of tumor-infiltrating OT-I, cells were stimulated overnight with 10 ng/mL of recombinant IL-2 (PeproTech, 402-ML) and 10^−8^ M ovalbumin epitope SIINFEKL in the presence of brefeldin A (Thermo Fisher Scientific, 00-4506-51). Afterwards, cells were stained for surface markers (see above), followed by fixation with BD Cytofix/Cytoperm (BD Biosciences, 51-2090KZ) and permeabilization with BD Perm/Wash (BD Biosciences, 51-2091KZ). Cells were stained with conjugated monoclonal antibodies directed at IL-2 (JES6-5H4, 561061, RRID:AB_395386) IFNγ (XMG1.2, 554411, RRID:AB_395375) from BD Biosciences, Granzyme B (NGZB, 12-8898-82, RRID:AB_10870787) and TNFα (MP6-XT22, 17-7321-82, RRID:AB_469508) from Thermo Fisher Scientific or Granzyme B (GB11, 560212, RRID:AB_11154033) from BD Biosciences.

Flow cytometry data were acquired on a CytoFLEX LX (Beckman Coulter Life Sciences) and analyzed using FlowJo software (v10.8.1, BD Biosciences, RRID:SCR_008520).

### Single-cell RNA sequencing

#### Cell preperation

Single-cell suspensions from tumors were obtained as described above (*Cell preparation*). Cells were labelled with TotalSeq-B anti-mouse hashtag antibodies and at the same time stained with an antibody mix used for sorting cell population of interest. 13 different TotalSeq-B antibodies were used to distinguish individuals (Hashtag antibody 1-13, BioLegend, 155831, 155833, 155835, 155837, 155839, 155841, 155843, 155845, 155847, 155849, 113901, 113903, 113905). At 4 degrees, cells were stained for Live/Dead for 20 minutes in 100 uL of PBS, then washed in PBS 10% FBS, then incubated with Fc receptor–blocking antibodies for 10 minutes. Without washing, hashing-sorting mix was added and incubation lasted for 30 min. LIVE/DEAD^−^TCRβ^+^CD8α^+^ cells were sorted from tumor-cell suspensions using a BD FACSAria III. After sorting, cell suspensions were filtered and pooled before being centrifuged for 10 min at 300g. Pellets were resuspended in RPMI 10% FBS and loaded on the Chromium Controller (10x Genomics) for single-cell RNA sequencing (scRNA-seq).

#### Library preparation

scRNA-seq libraries were prepared using the Chromium Single Cell 3’ v3.1 Reagent Kit (10x Genomics) according to manufacturer’s user guide CG000317. Libraries were loaded to an Illumina Novaseq (Brightcore platform) and Cell Ranger 9.0.1 (10x Genomics) function count were used to demultiplex the sequencing data, generate gene-barcode matrices, and align the sequences to the mm10 genome. Resulted matrices were used to generate a single cell seurat object using Seurat 5.1.0 R package.

#### Demultiplexing

Data containing information about oligo-tagged antibodies and cell barcodes were added as an independent assay in the Seurat object and were normalized with the NormalizeData function from the Seurat package using the “CLR” normalization method which applies a centered log ratio transformation. The HTODemux function was used to reassign individual cells to their original mouse sample. Negative cells with no oligo-tagged antibody detected were removed from downstream analysis.

#### Filtering

RNA data were filtered by mouse sample with the following thresholds: number of genes expressed > 700 (between 700 and 4500 depending on the mouse sample), fraction of mitochondrial content < 5%. The new data from the filtered Seurat object were normalized with NormalizeData() and scaled with ScaleData() functions from Seurat package. With the following variables to regress: ncount_RNA, mitochondrial and ribosomal content.

Final-filtered Seurat object was generated using SCTransform() function in which the top 5000 variable genes were identified. These variable genes were used to perform and select top 20 principal components analysis (PCA). These components were then used to build a Uniform Manifold Approximation and Projection (UMAP) embedding of the 11025 cells. Unsupervised clustering of the cells was performed with the Louvain algorithm to determine the different cell clusters (resolution = 0.8).

#### Differentially expressed genes (DEG) analysis

Genes differentially expressed among resulted clusters were determined using the FindAllMarkers function of the Seurat package. Genes with an adjusted p-value of 0.05 and with a logFoldChange absolute value greater than 0.25 were retained and represented as a volcano plot using the EnhancedVolcano 1.22.0 R package.

#### Trajectory analysis

The Slingshot 2.12.0 R package was used to perform trajectory analysis on the filtered Seurat object and pseudo-time of lineages was calculated with the following parameters: extend = ‘n’, start.clus =“Tcm1”, approx_points =F,end.clus=“Tex”.

## Acknowledgements

We especially thank Oberdan Leo and Muriel Moser for their support all along this work, for stimulating discussions and for careful revision of the manuscript. We also thank the entire team at the animal care facility, and Isabelle Decot and Véronique Dissy for administrative and technical support. This study was supported by the Fonds National de la Recherche Scientifique (FRS-FNRS, Belgium), the European Regional Development Fund (ERDF) of the Walloon Region (WAL-IMAGIN portfolio, SYST-IMM), the Fonds Cancer Hainaut, the Fonds Genicot and Fonds Jaumote-Demoulin. HD, FB and ST are recipient of FNRS-Télévie grant. HD has been supported by the Rose et Jean Hoguet Fund. MLM and SD are research fellows, and FA and SG are senior research associate and research director of the FRS-FNRS.

