## supplemental figure for "TCR Signal Strength and Eomes Coordinate CD8^+^ T Cell Fate and Antitumor Immunity in Hepatocellular Carcinoma"

Supplemental figures 1 - 14

**Supplementary figure 1 – Hydrodynamic plasmid injection induces an immune response**
**(A)** Plasmids injected into mice to induce hepatocarcinoma. **(B)** Representative FACS plots and frequencies of OVA-pentamer$^+$ cells among CD8a$^+$ T cells from ctrl and OVA-HCC mice 5 weeks after plasmid injection. Data are shown as median with interquartile range and are representative of 5 experiments (n= 15 to 35 mice per group), each dot represents a single mouse. Statistical significance was determined using the Mann-Whitney test, ****p<0.0001.

**Supplementary figure 2 – Transfer of OT-I cells induce a transient tumor regression**
**(A)** Schematic representation of the experimental design used to study T cell responses to hepatocarcinoma (upper panel) and combinations of plasmids injected in OVA-HCC and OVA-liver settings. **(B)** Bioluminescence imaging (BLI) quantification of tumor burden before (day 0) and following OT-I cells transfer during 4 weeks. **(C)** Pictures of the liver in each mice group. **(D)** Liver weight in each mice group. **(E, H, K)** Absolute number of OT-I cells (E), Tex-like cells (H), and terminally exhausted T cells (K) in OVA$^{HCC}$ and OVA$^{Liver}$ mice. **(F, G, I, J)** Representative FACS profiles (F-I) and frequencies (G-J) of TOX$^+$PD1$^+$ Tex-like cells among OT-I CD8$^+$ T cells and TCF1$^-$TIM3$^+$ terminally exhausted T cells (Ttex) among PD1$^+$ TOX$^+$ cells. **(L, M)** Representative FACS plots (L) and frequencies (M) of of IFNγ$^+$TNFα$^+$ OT-I cells in each group. Data are shown as median with interquartile range and are pooled from 3 to 6 experiments (E-K) or representative of 2 experiments (L, M) (n= 4-26 mice per group), each dot represents a single mouse. Statistical significance was determined using the Mann-Whitney test, ns = non-significant, *p<0.05, ***p<0.0001.

**Supplementary figure 3 – High affinity tumor-specific T cells promote Trm and Teff differentiation in the liver**
**(A)** Representative images of livers from N4, Q4, T4 and Ctrl (without tumor inducing plasmids) groups. **(B)** t-SNE analysis of N4 and Q4 groups showing expression of selected markers. **(C)** representative flow cytometry plots of Granzyme B (GranzB), interferon gamma (IFNγ), tumor necrosis factor alpha (TNFα), and interleukin-2 (IL-2). Representative data from one experiment with n = 5 mice per group.

**Supplementary figure 4 – High affinity tumor-specific T cells promote Trm and Teff differentiation in the spleen**
Frequencies, absolute numbers, and representative FACS plots of TCF1$^-$CX3CR1$^+$ (Teff-like), CXCR6$^+$CD49a$^+$ (Trm-like) and PD1$^+$TOX$^+$ (Tex-like) cells gated on CD8$^+$ T cells in N4 and Q4 groups. Data are shown as median with interquartile range and are representative of 2 experiments (n= 11 to 12 mice per group), each dot represents a single mouse. Statistical significance was determined using the Mann-Whitney test, ns = non-significant, *p<0.05, **p<0.01, **p<0.001.

**Supplementary figure 5 – Eomes positively correlates with TCF1 expression and negatively correlates with CX3CR1 and CD49a expression**
**(A-C)** Representative FACS plots (left panels) and median fluorescence of CD49a, CX3CR1 and TCF1 in Eomes+ and Eomes- cells (right panels) from the N4 and Q4 groups. Data are shown as median with interquartile range and are representative of 2 experiments (n= 6 mice per group), each dot represents a single mouse. Statistical significance was determined using the Multiple Mann-Whitney test, ns = non-significant, ns = non-significant, **p<0.01.

**Supplementary figure 6 – Eomes depletion improves tumor control**
Bioluminescence imaging (BLI) quantification of tumor burden before (day 0) and following OT-I cells transfer during 3 weeks in WT and Eomes-KO groups (individual mice from Fig. 2D).

Data are shown as median with interquartile range and are representative of 3 experiments with n= 12-13. Statistical significance was determined using the Mann-Whitney test, ns = non-significant, **p<0.01. Mann-Whitney test was corrected for multiple comparisons using Holm-Šídák method.

**Supplementary figure 7 – Gene expression and signature profiling for cell population annotation**

**(A)** Violin plots showing the expression of different genes across annotated cell populations.
**(B)** Features plots showing expression levels of selected genes across annotated cell populations.
**(C)** Violin plots showing expression levels of literature-derived gene signatures across annotated cell populations.

**Supplementary figure 8 – Eomes depletion enhances the expression of tissue-residency-associated genes**

**(A-E)** Volcano plot comparing gene expression in Tcm2 (A), Tcm3 (B), Tpex (C), Tem (D), Tex-eff (E) populations from WT and Eomes-KO OT-I cells isolated from HCC livers. Each dot represents one gene. Red dots correspond to p value < 0,01 and fold change > 0,5. **(F)** Violin plots showing expression levels of TRM_KODA signature genes across annotated cell populations, split into WT and Eomes-KO groups.

**Supplementary figure 9 – Eomes depletion limits exhausted T cells differentiation and increases the proportion of Trm cells.**

**(A-C)** Representative FACS plots at day 7 and day 21 showing $CXCR6^+CD49a^+$ (Trm-like) (A), $PD1^+TOX^+$ (Tex-like) (B) and $TCF1^-CX3CR1^+$ (C) populations gated on $CD8^+$ OT-I cells from WT and Eomes-KO groups. **(D)** Histogram showing Tox expression among $TCF1^-CX3CR1^+$ cells in each group. **(E)** Frequency of $Tox^+$ (Tex-eff) and $Tox^-$ (Teff) cells among TCF1-CX3CR1+ cells in each condition.

**Supplementary figure 10 – Eomes deficiency reduces the proportion of KLRG1-expressing cells**

Representative FACS plots and frequencies of KLRG1 and CX3CR1 populations among OT-I cells in WT and Eomes-KO groups a day 7 and 21. Data are shown as median with interquartile range and are representative of 1 experiment (n= 5 to 4 mice per group), each dot represents a single mouse. Statistical significance was determined using the Multiple Mann-Whitney test, ns = non-significant, ns = non-significant, *p<0.05, **p<0.01, ***p<0.001, ****p<0.0001.

**Supplementary figure 11 – Eomes depletion increases the frequency of cytokine-producing cells**

**(A)** t-SNE analysis of $CD8^+$ OT-I cells at day 21 in WT and Eomes-KO mice, showing expression of selected markers. **(B, C)** t-SNE (B) and Pie charts (C) indicate the proportion of cells secreting 0, 1, 2 or 3 cytokines in each group. n = 4 mice per group.

**Supplementary figure 12 – Eomes deficiency enhances Trm cell development in a low-affinity HCC model**

t-SNE analysis of Q4 WT and Q4 Eomes-KO groups, showing expression of selected markers. Data pooled from 4 mice per group.

**Supplementary figure 13 – OT-I Eomes transgenic cells express physiological levels of Eomes and display reduced proliferative capacity**

**(A)** Histogram of the expression of Eomes in $CD8^+$ T cells from liver and spleen in B6, OT-I $Rag1^{-/-}$ and OT-I $Rag1^{-/-}$ $Eomes^{Tg/Tg}$ mice. **(B)** Representative dot plots of $PD1^+TOX^+$ among CD8+ T cell in OT-I $Rag^-/^-$

and OT-I Rag-/- $Eomes^{Tg/Tg}$ mice. **(C)** In vitro proliferation of WT, Eomes-KO and $Eomes^{Tg/Tg}$ OT-I cells in response to OVA. Representative data from one experiment with n = 2 mice per group.

**Supplementary figure 14 – transgenic Eomes expression promotes Tex differentiation while reducing Teff and Trm cell development**

**(A)** Fold change of BLI quantification of tumor burden at day 7 in recipient mice of WT and $Eomes^{Tg/Tg}$ OT-I cells. **(B)** t-TSNE analysis of WT and $Eomes^{Tg/Tg}$ groups, showing expression of selected markers. **(C-E)** Representative flow cytometry plots of CXCR6 CD49a, TCF1 CX3CR1, TOX PD1 among $CD8^+$ T cells in each group. Data are shown as median with interquartile range and are pooled from 3 independent experiments (n= 23 to 24 mice per group), each dot represents a single mouse. Statistical significance was determined using the Multiple Mann-Whitney test, ns = non-significant.

**Supplementary figure 1**

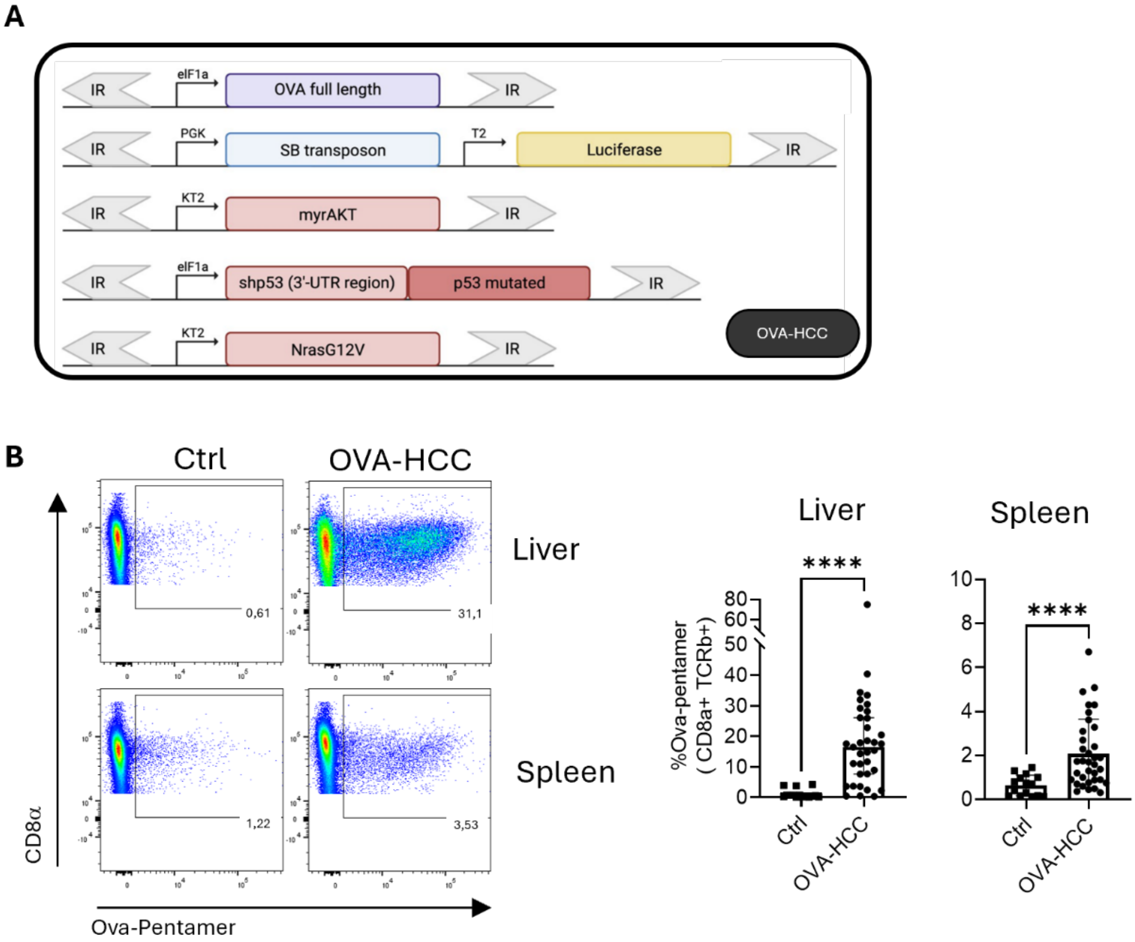

**Supplementary figure 2**

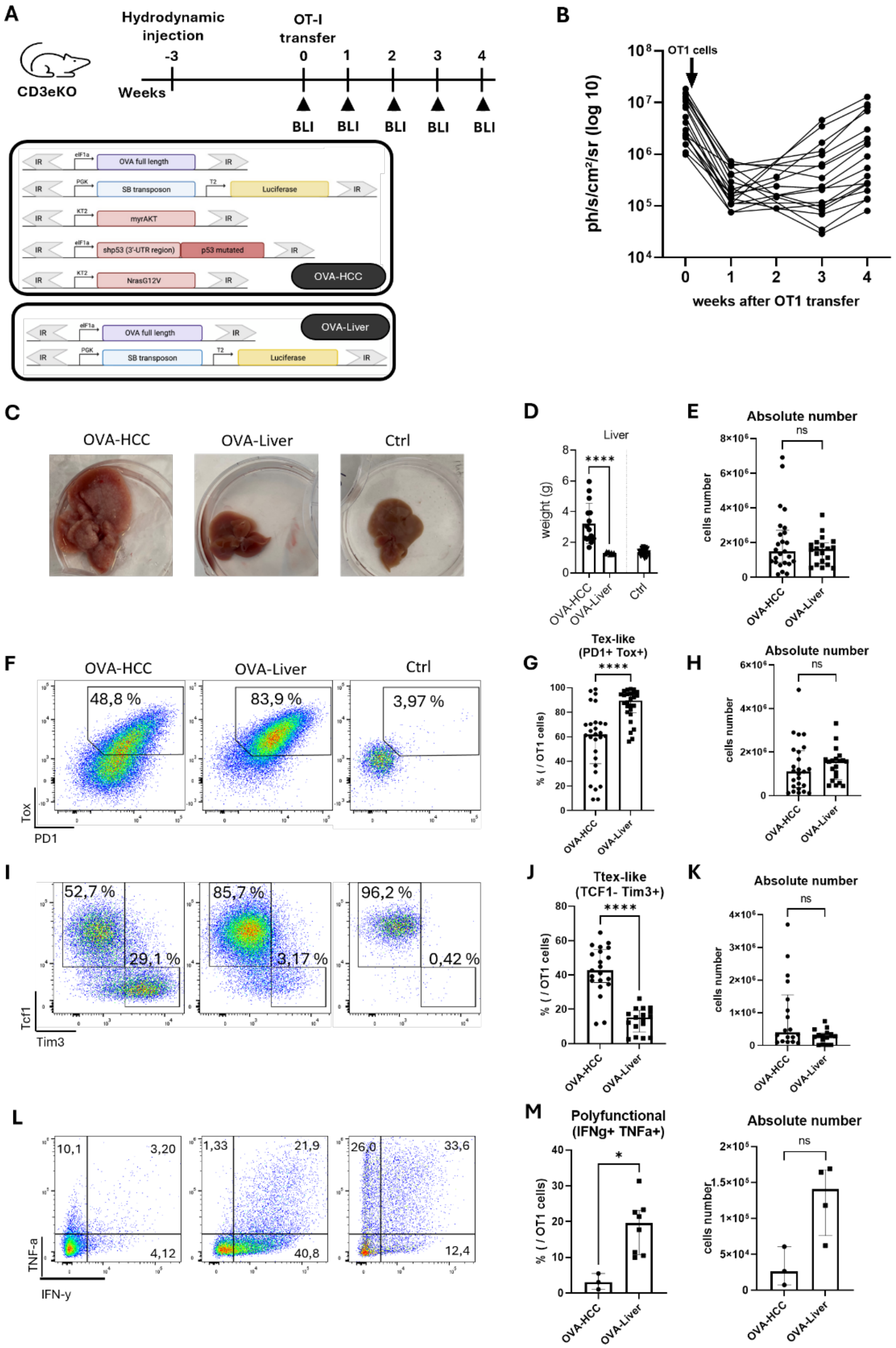

**Supplementary figure 3**

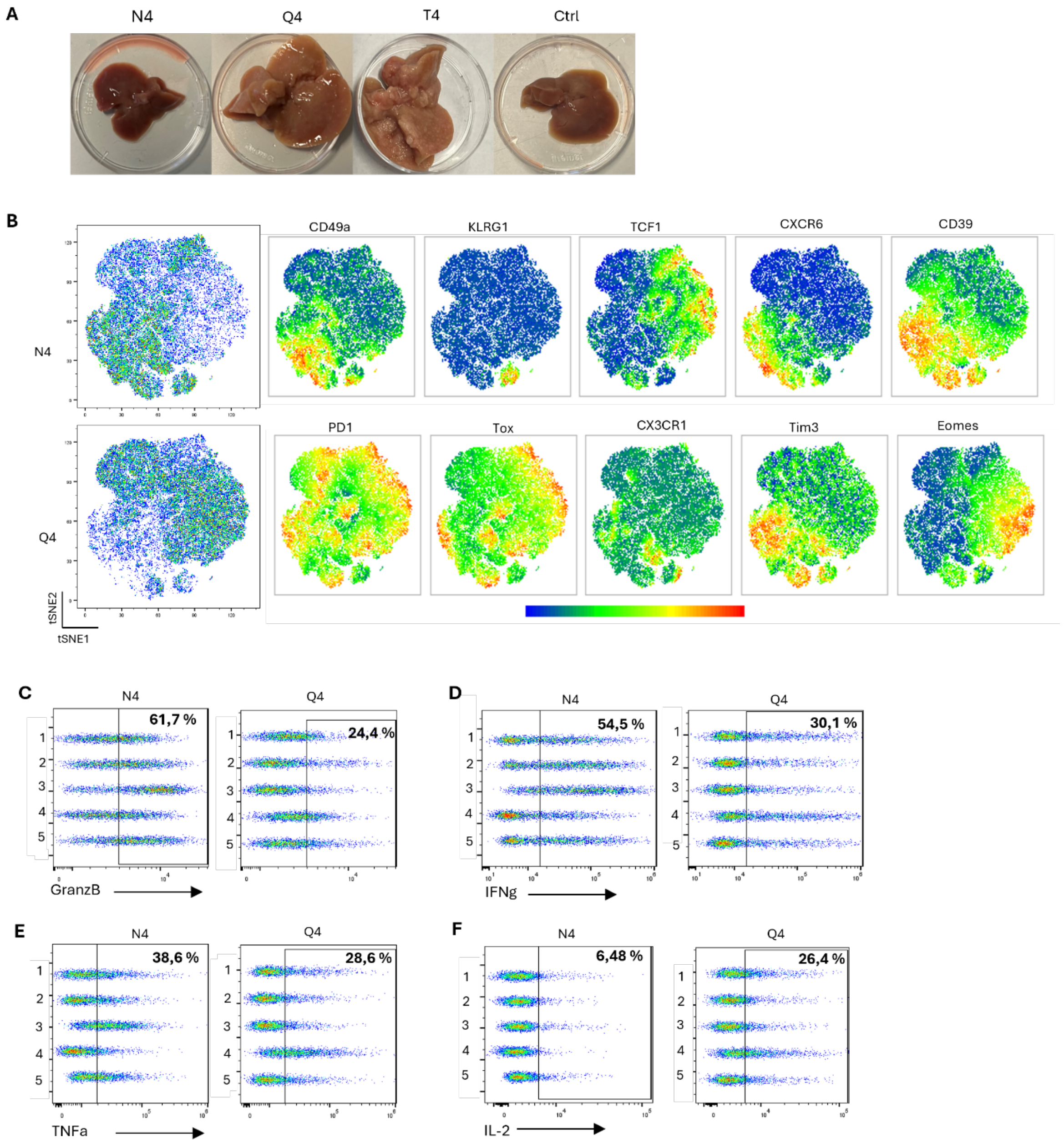

**Supplementary figure 4**

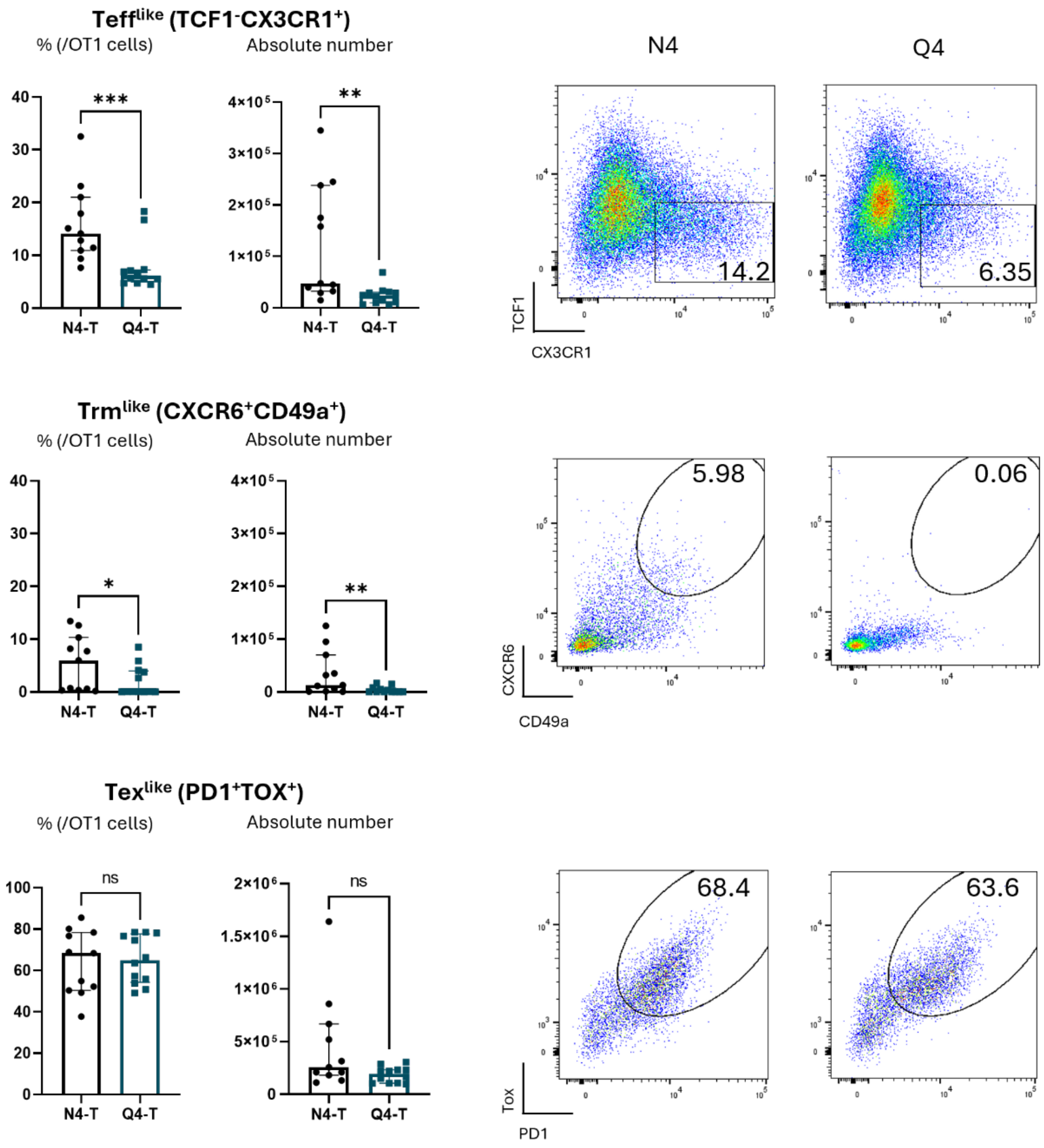

**Supplementary figure 5**

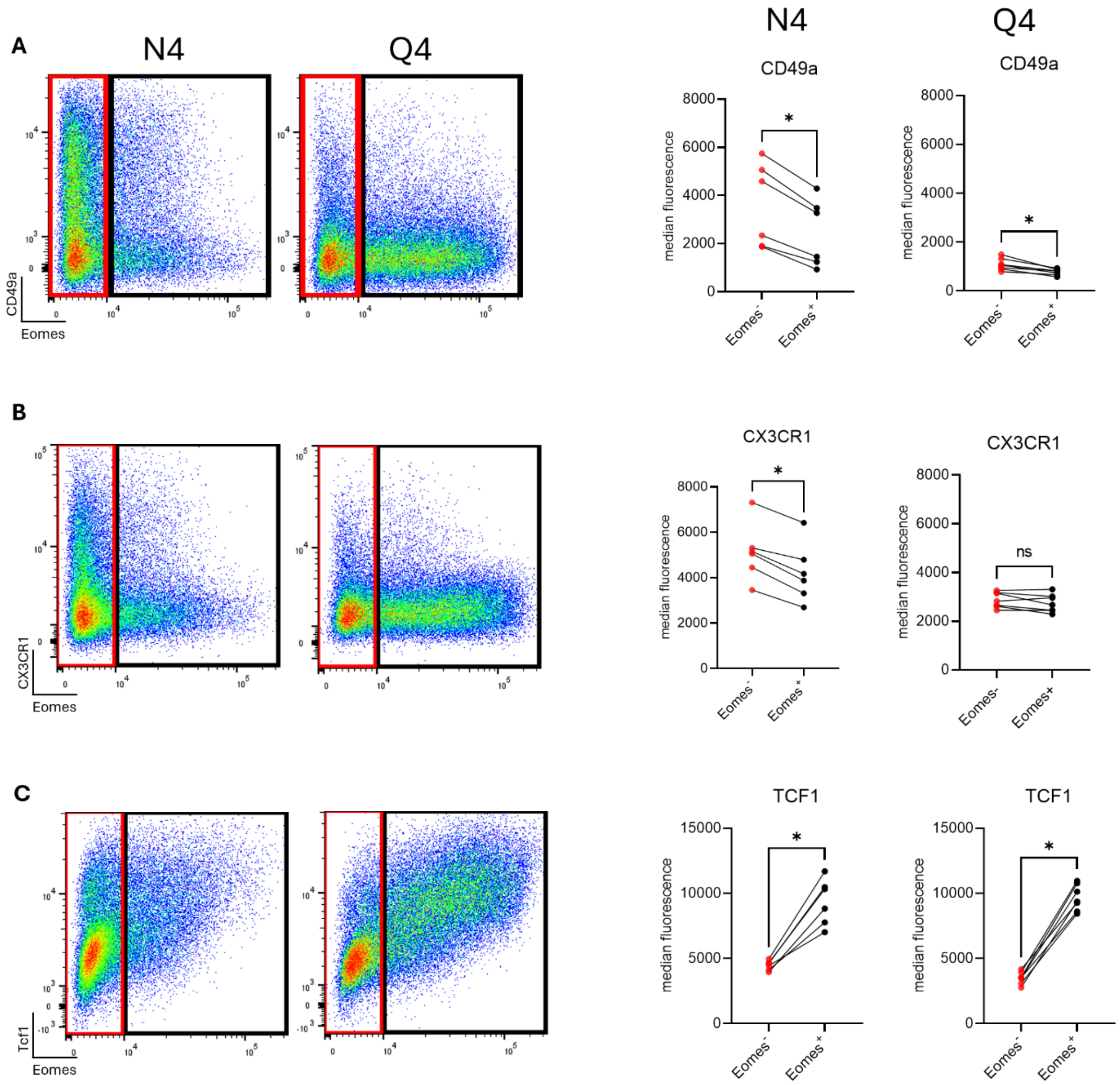

**Supplementary figure 6**

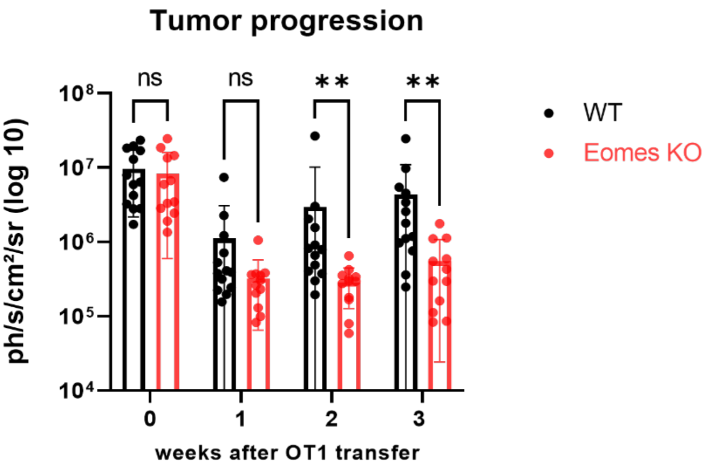

**Supplementary figure 7**

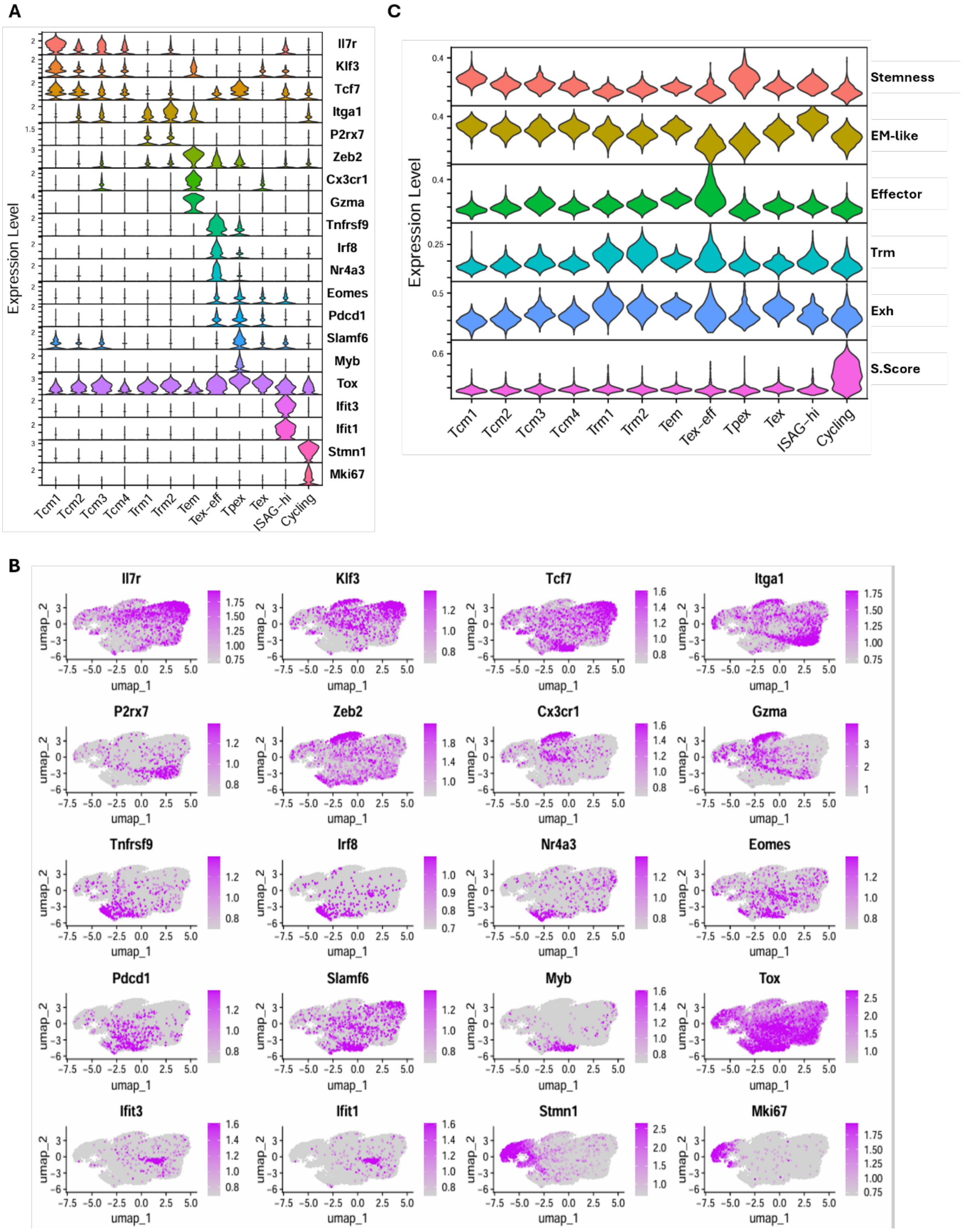

**Supplementary figure 8**

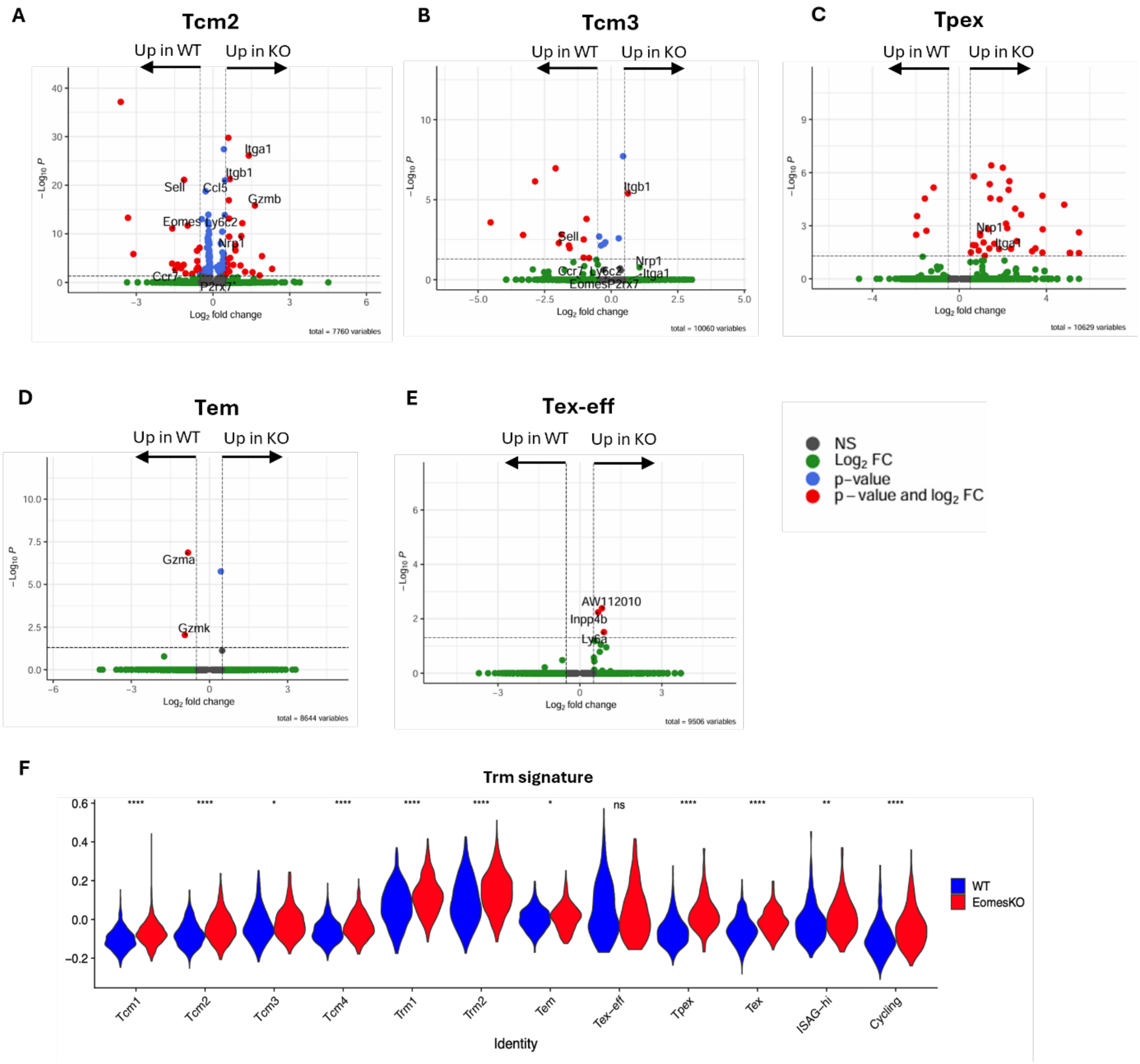

**Supplementary figure 9**

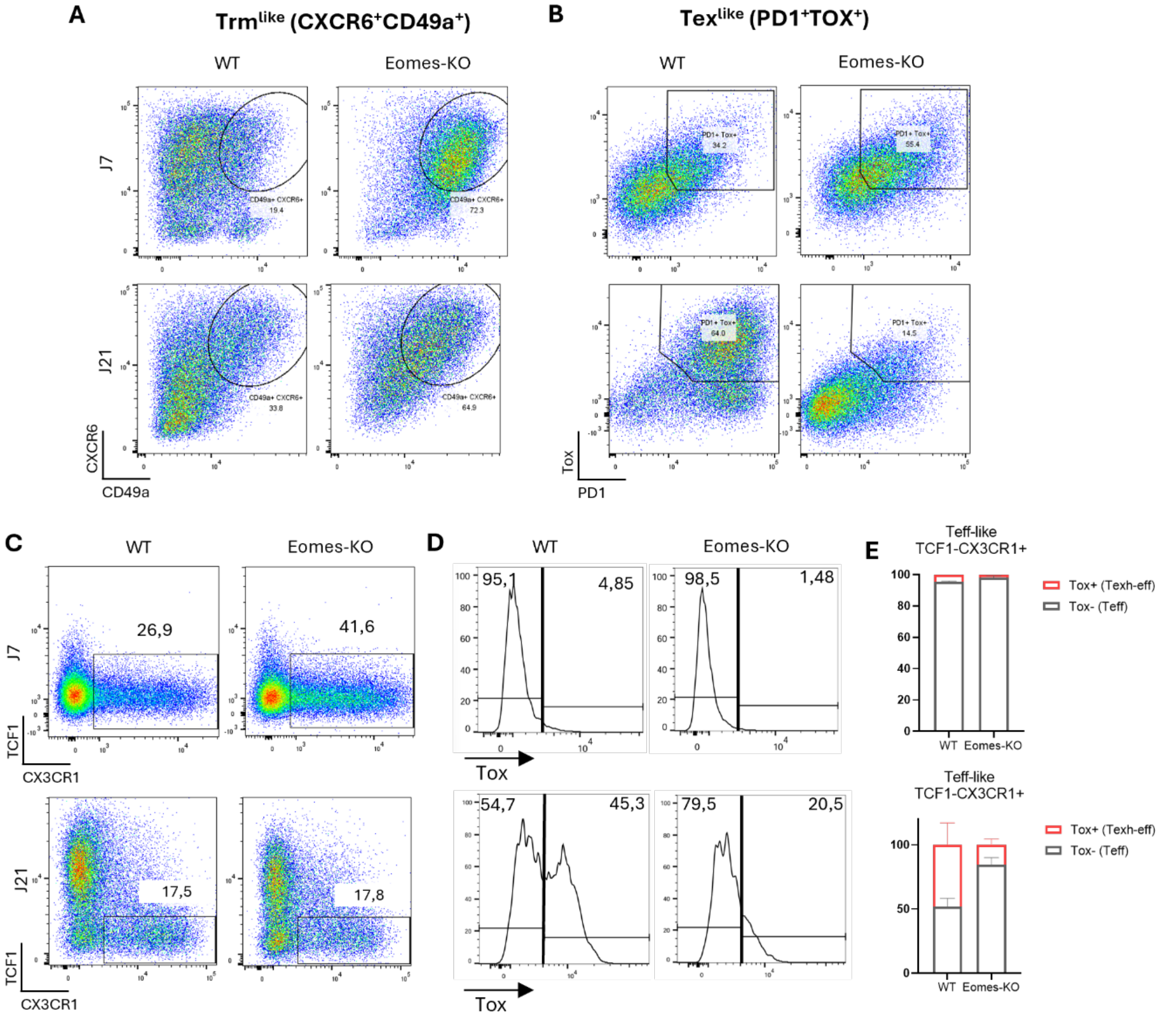

**Supplementary figure 10**

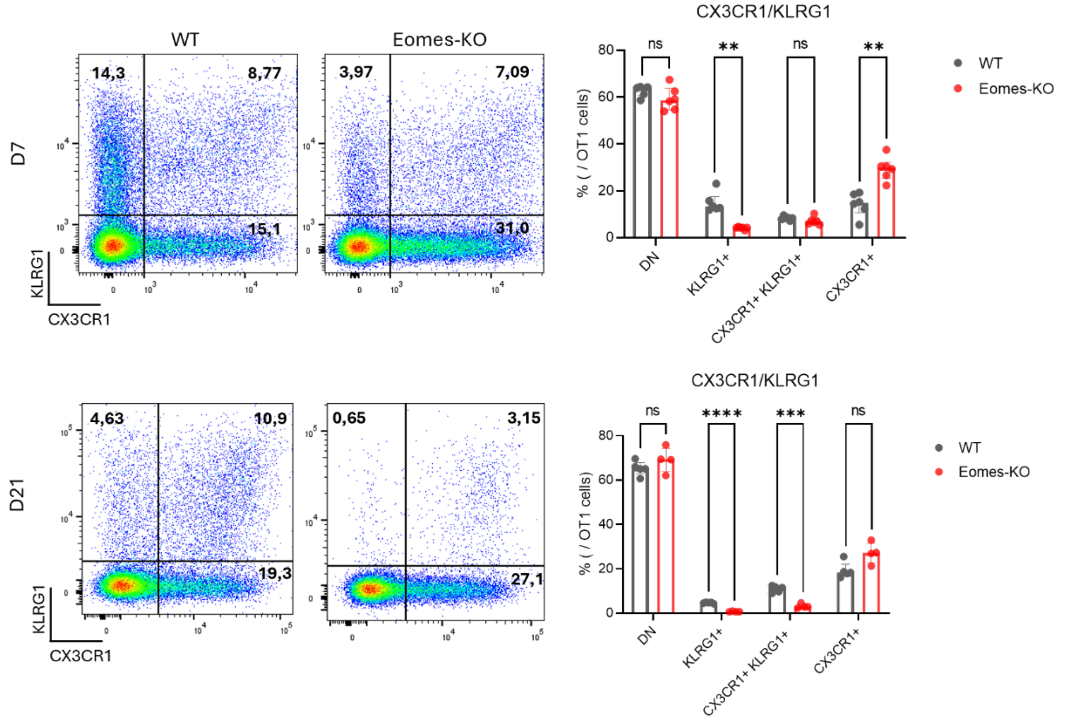

**Supplementary figure 11**

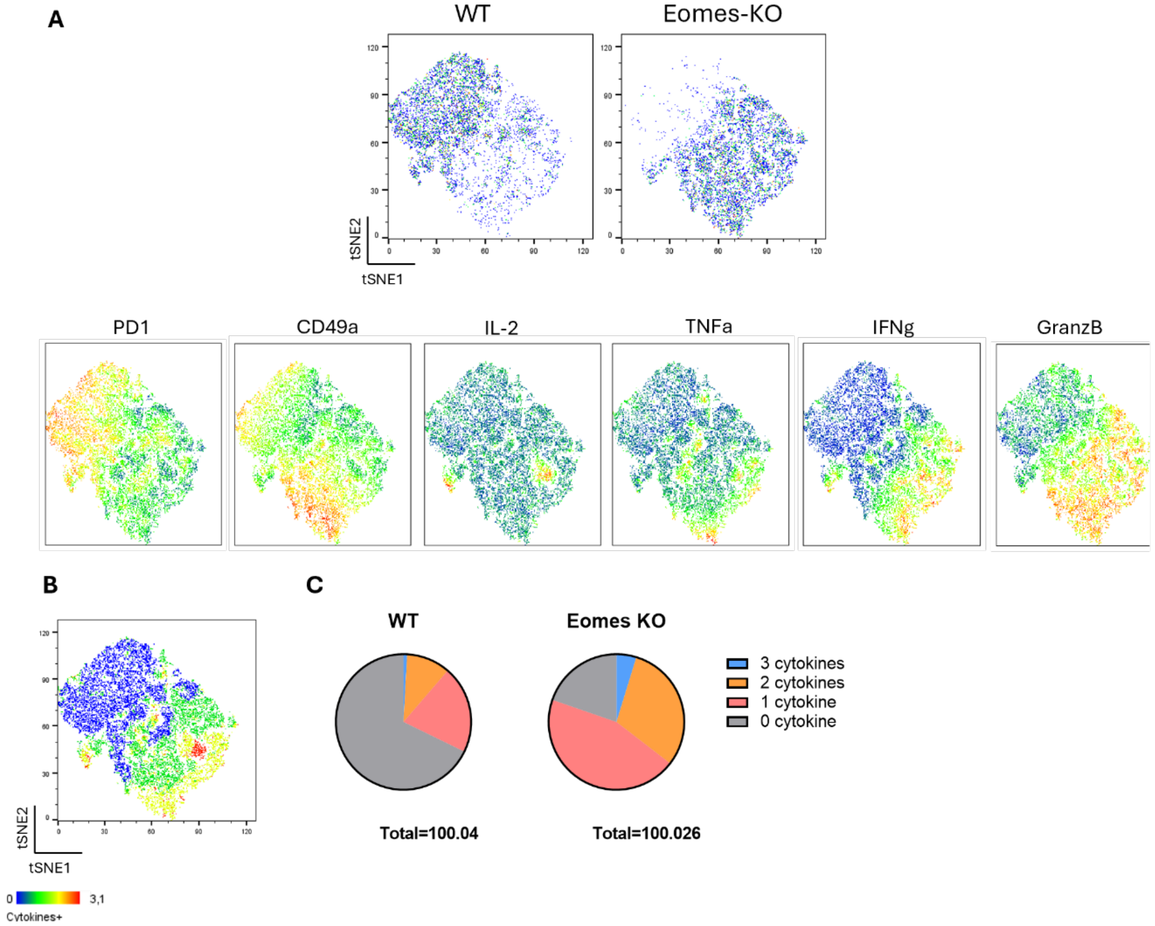

**Supplementary figure 12**

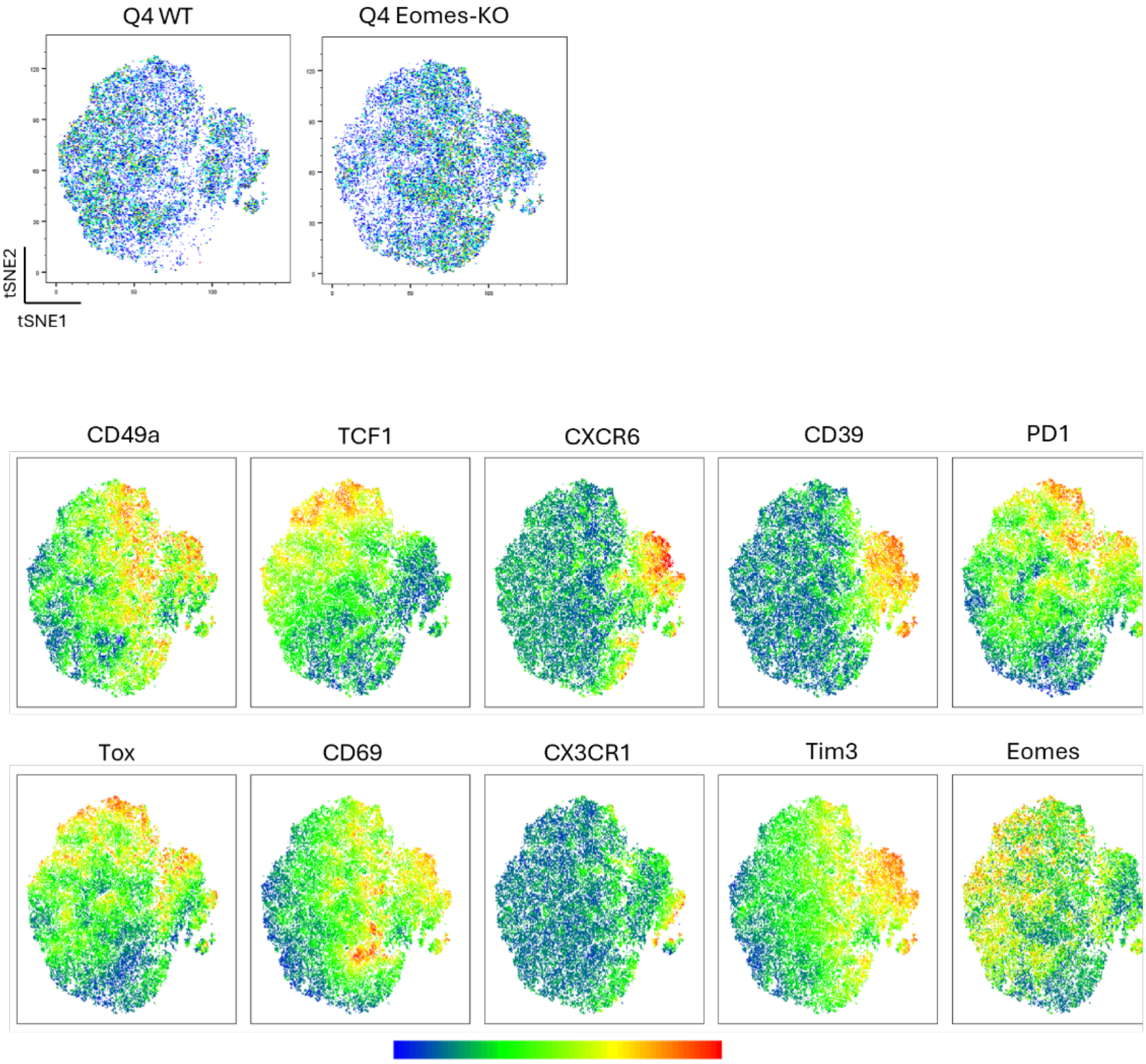

### Supplementary figure 13

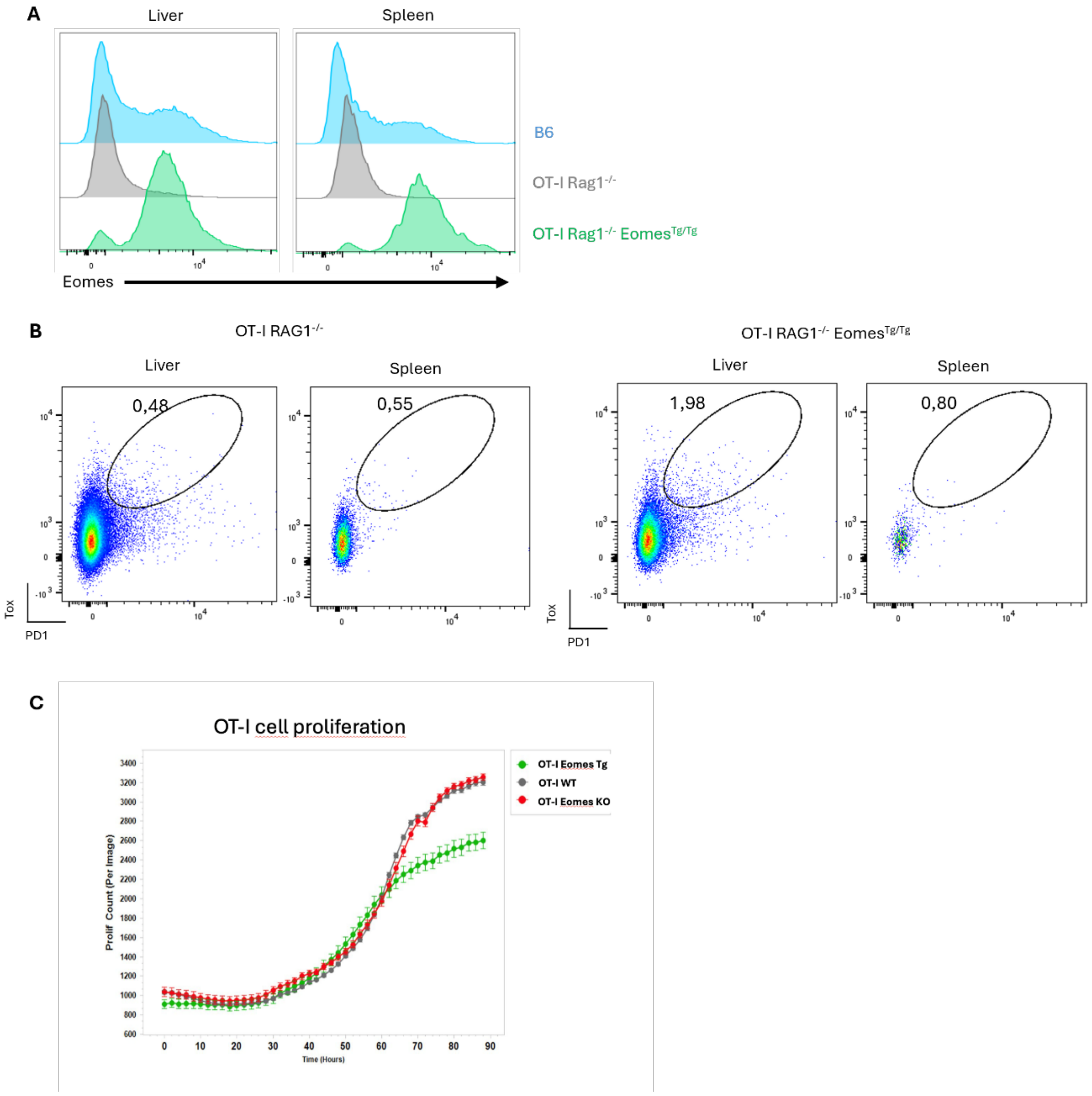

**Supplementary figure 14**

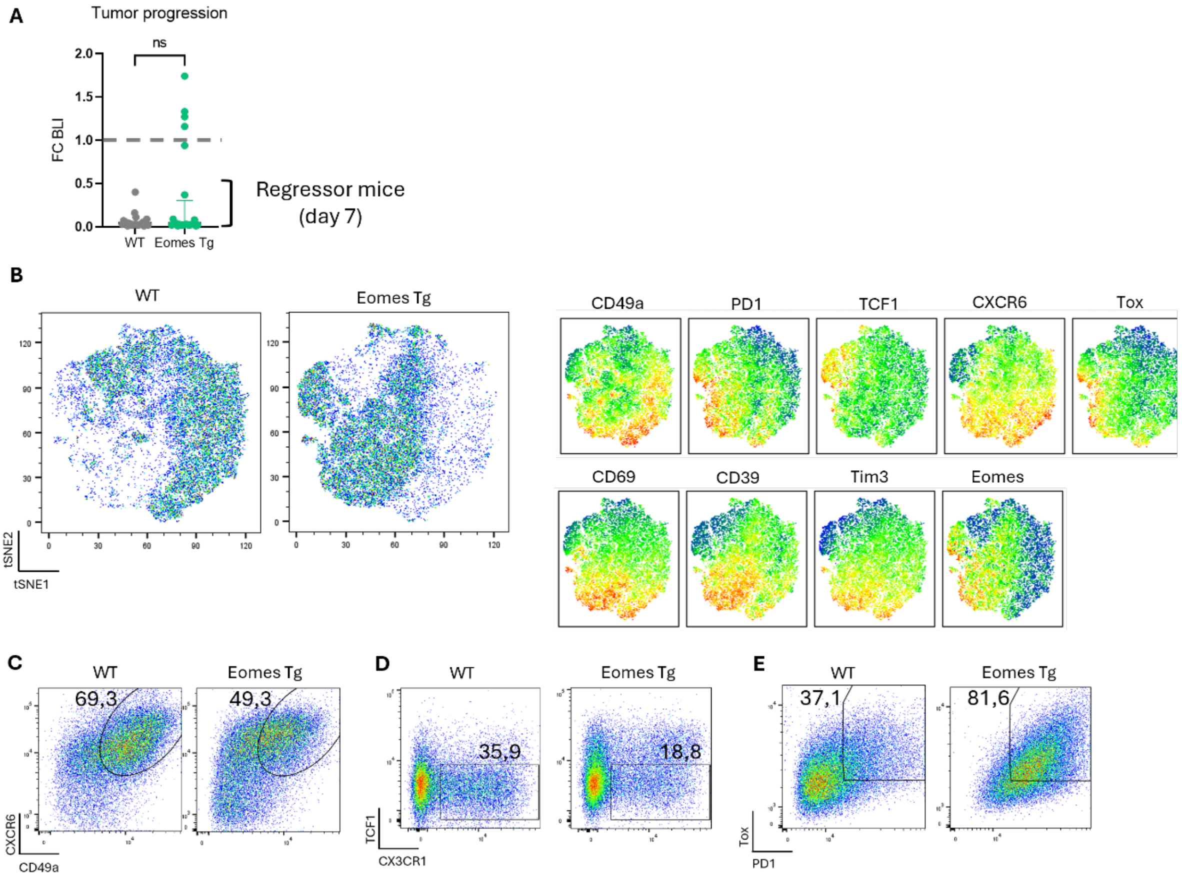
